# Dependence-Aware Conformal Prediction for 3D Uncertainty Quantification in Molecular Docking

**DOI:** 10.64898/2026.11.20.768203

**Authors:** Guanyu Lyu, Haochang Hao, Yihang Wang, Lu Cheng

## Abstract

Molecular docking plays a critical role in drug discovery by predicting three-dimensional (3D) binding poses of small molecules within proteins. However, docking predictions can contain substantial positional errors, making reliable uncertainty quantification essential. Conformal prediction (CP) provides a principled framework for uncertainty quantification, yet constructing compact 3D prediction regions with reliable joint coverage remains challenging due to heterogeneous atomic errors, dependence among spatial coordinates, and distribution shifts. We develop a dependence-aware CP framework for molecular docking that leverages a shared graph neural network (GNN) to estimate adaptive, asymmetric uncertainty intervals using local atomic and global molecular information. Copula-based joint calibration accounts for dependence among coordinate-wise nonconformity scores to construct compact 3D prediction boxes. We further introduce density ratio estimation (DRE) to adapt calibration under distribution shift. Experiments across three docking methods demonstrate that our approach reduces median prediction-box volumes by 37.40%, 95.00%, and 39.16% for AutoDock Vina, Protenix, and MedusaGraph, respectively, compared with a state-of-the-art CP baseline, while achieving empirical joint coverage close to the 90% target. Under the Molecular Group Split and controlled distribution shifts, DRE further reduces coverage deviations, highlighting the potential of dependence-aware and shift-adaptive calibration for reliable 3D molecular docking uncertainty quantification.

## 1 Introduction

Predicting how small molecules interact with proteins is fundamental to computational biology and drug discovery. Molecular docking predicts the binding pose of a small molecule (ligand) within a target protein, represented by the three-dimensional (3D) coordinates of its atoms (Morris & Lim-Wilby, 2008; Paggi et al., 2024). Accurate predictions help researchers understand molecular interactions and identify promising drug candidates. Recent docking methods can be broadly categorized into three groups: search-and-scoring methods such as AutoDock Vina (Trott & Olson, 2010; Eberhardt et al., 2021) and GNINA (McNutt et al., 2021), deep learning-based docking methods such as MedusaGraph (Jiang et al., 2022) and DynamicBind (Lu et al., 2024), and protein–ligand co-folding methods such as Protenix (ByteDance AML AI4Science Team et al., 2025) and AlphaFold 3 (Abramson et al., 2024). However, predicted structures rarely indicate the reliability of individual atomic positions. Even physically plausible binding poses may contain substantial positional errors, motivating the need for reliable uncertainty quantification. A fundamental question is:

### How can we construct reliable and informative 3D uncertainty regions for individual atoms in molecular docking predictions?

Each atom has three spatial coordinates whose prediction errors may be statistically dependent, while error magnitudes vary across atoms and protein–ligand complexes. Effective uncertainty quantification must therefore account for both coordinate dependence and heterogeneous errors, producing compact prediction regions with reliable joint coverage.

Conformal prediction (CP) provides a principled, distribution-free framework for uncertainty quantification, offering finite-sample marginal coverage guarantees under the exchangeability assumption (Vovk et al., 2005; Angelopoulos & Bates, 2023; Zhou et al., 2025; Angelopoulos et al., 2024). By leveraging calibration data and nonconformity scores, CP constructs prediction sets with guaranteed coverage without requiring restrictive assumptions about the underlying predictive model or error distribution. Recent studies have explored CP for molecular predictions (Alvarsson et al., 2021; Nigam et al., 2021), including graph neural network (GNN)-based atom-level uncertainty estimation for docking (Hao et al., 2026). However, constructing efficient 3D prediction regions remains challenging. Independently calibrated coordinate intervals do not guarantee joint coverage, whereas conservative coordinate-wise corrections can yield excessively large regions. Although copula-based CP has been investigated for multivariate prediction (Messoudi et al., 2021; Sun & Yu, 2024), capturing coordinate dependence alongside heterogeneous atomic errors remains underexplored in molecular docking. Distribution shift introduces an additional difficulty. Docking models may encounter proteins and ligands that differ from the calibration data, potentially altering prediction-error distributions and compromising coverage. Weighted CP can address certain forms of covariate shift through density-ratio-based calibration (Tibshirani et al., 2019; Barber et al., 2023). However, adapting this approach to molecular docking requires estimating distribution differences across complex protein–ligand systems while accounting for dependence among spatial coordinates.

To address these challenges, we propose a dependence-aware CP framework for 3D atom-level uncertainty quantification in molecular docking. A shared GNN with GATv2 message passing (Brody et al., 2022) uses local atomic features and global molecular information to estimate asymmetric, atom-specific uncertainty widths. We then employ empirical, Gaussian, and R-vine copulas to model dependence among coordinate-wise nonconformity scores, constructing compact 3D prediction boxes while targeting joint coverage. Finally, density ratio estimation (DRE) based on learned molecular representations enables adaptive calibration under distribution shift. We evaluate our framework on three representative docking methods: AutoDock Vina, Protenix, and MedusaGraph under independent and identically distributed (IID) random splits and distribution shifts. Our contributions are:

- We develop a 3D atom-level CP framework that leverages GNN-based uncertainty estimation and copula-based joint calibration to account for heterogeneous prediction errors and coordinate dependence.
- We incorporate density-ratio-based calibration using learned molecular representations to improve joint coverage under distribution shift and investigate when this adaptation is effective.
- Experiments demonstrate that empirical copula calibration reduces median prediction-box volumes up to 95.00%, compared with the state of the art, while maintaining empirical joint coverage near 90%. DRE further reduces coverage deviations under distribution shifts.

## 2 Related Works

### Uncertainty Quantification for Molecular Docking

Reliable uncertainty quantification is important in molecular docking, where predicted protein–ligand binding poses may contain substantial positional errors. Existing approaches have investigated different forms of confidence and uncertainty estimation. GNINA (McNutt et al., 2021) employs an ensemble of convolutional neural networks for docking pose scoring, while DiffDock (Corso et al., 2023) uses a learned confidence model to assess predicted binding poses. Beyond pose-level confidence, ensemble-based and Bayesian methods have been investigated for uncertainty quantification in protein–ligand binding affinity prediction (Rayka et al., 2024; Rayka & Naghavi, 2025). Stochastic modeling has also been explored to characterize uncertainty in receptor–ligand binding-site prediction (Chen et al., 2024). However, these approaches generally do not provide calibrated three-dimensional (3D) uncertainty regions for individual ligand atoms. More recently, ConfDock (Hao et al., 2026) has explored GNN-based conformal prediction for adaptive, coordinate-wise atom-level uncertainty estimation. Our work focuses on dependence-aware joint calibration, which provides a statistically principled way to calibrate empirical joint coverage without assuming coordinate-wise independence and yields compact 3D prediction regions.

### Conformal Prediction under Dependence and Distribution Shift

CP provides distribution-free marginal coverage guarantees under exchangeability (Vovk et al., 2005; Papadopoulos et al., 2002; Lei et al., 2018; Zhou et al., 2025). Conformalized Quantile Regression (CQR) (Romano et al., 2019) further enables adaptive prediction intervals for heterogeneous errors. For multivariate predictions, however, coordinate-wise coverage does not guarantee simultaneous coverage of all coordinates. Copula-based CP addresses this challenge by modeling dependence among nonconformity scores, as explored in multi-target regression (Messoudi et al., 2021) and multi-step time-series forecasting (Sun & Yu, 2024). Distribution shift introduces another challenge, as violations of exchangeability can compromise coverage. Weighted CP (Tibshirani et al., 2019) and extensions beyond exchangeability (Barber et al., 2023) provide foundations for addressing such shifts. DRE, including domain-classification approaches (Bickel et al., 2009; Sugiyama et al., 2012), estimates distribution differences to reweight calibration samples. However, errors in estimated density ratios can weaken coverage guarantees (Bhattacharyya & Barber, 2026). Our work investigates coordinate dependence and distribution shift in molecular docking through copula-based joint calibration and DRE using GNN-derived molecular representations. Specifically, we apply estimated density-ratio weights during joint copula calibration while retaining unweighted marginal transformations, allowing joint calibration thresholds to adapt to distribution shifts.

## 3 Preliminaries and Problem Setup

**Conformal Prediction** CP constructs prediction sets with finite-sample marginal coverage guarantees under exchangeability (Vovk et al., 2005; Papadopoulos et al., 2002; Lei et al., 2018). Given a trained prediction model and a held-out calibration set 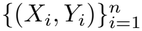, where *X*_*i*_ denotes the input and *Y*_*i*_ its corresponding outcome, split CP uses nonconformity scores *s*(*X*_*i*_, *Y*_*i*_) to construct

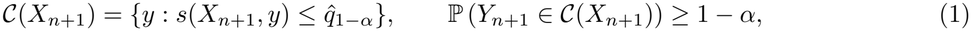

where *α* ∈ (0, 1) is the target miscoverage level, *X*_*n*+1_ and *Y*_*n*+1_ denote the test input and outcome, and 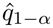 is the ⌈ (*n* + 1)(1− *α*) ⌉ -th smallest calibration score (taken as +∞ if the index exceeds *n*). CQR further enables adaptive and asymmetric prediction intervals (Romano et al., 2019). However, when predicting multiple dependent quantities, calibrating each coordinate independently does not ensure their simultaneous coverage. Additional details on CP and exchangeability are provided in Appendix B.

### Copula-Based Dependence Modeling

Copulas characterize statistical dependence among random variables separately from their marginal distributions. By Sklar’s theorem (Sklar, 1959; Nelsen, 2006), the joint cumulative distribution function (CDF) of three coordinate-wise nonconformity scores (*S*_*x*_, *S*_*y*_, *S*_*z*_) is

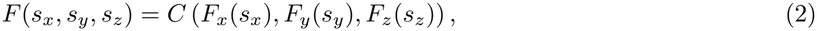

*F* (*s*_*x*_, *s*_*y*_, *s*_*z*_) = *C* (*F*_*x*_(*s*_*x*_), *F*_*y*_(*s*_*y*_), *F*_*z*_(*s*_*z*_)), (2) where *S*_*a*_ is the nonconformity score for coordinate axis *a*∈ *{x, y, z}, s*_*a*_ is a possible value of *S*_*a*_, *F*_*a*_(*s*_*a*_) = ℙ(*S*_*a*_ ≤*s*_*a*_) is its marginal CDF, *F* is the joint CDF, and *C* : [0, 1]^3^→ [0, 1] is a copula capturing dependence among the three scores. We consider empirical, Gaussian, and R-vine copulas, with details provided in Appendix E.

### Problem Formulation

Molecular docking predicts the 3D binding pose of a ligand within a target protein. We represent each protein–ligand complex as a molecular graph and denote its set of ligand atoms by *V*. For each atom *v* ∈ V, let **ŷ**_*v*_ = (*ŷ*_*v,x*_, *ŷ*_*v,y*_, *ŷ*_*v,z*_) ∈ ℝ^3^ and **y** = (*y*_*v,x*_, *y*_*v,y*_, *y*_*v,z*_) ∈ ℝ^3^ denote its predicted and ground-truth coordinates,respectively. Given the molecular graph, predicted coordinates, and a calibration dataset, our goal is to construct a compact 3D prediction box B = *I*_*v,x*_ × *I*_*v,y*_ × *I*_*v,z*_,, where *I*_*v,a*_ ⊆ ℝ is the prediction interval for atom *v* along coordinate axis *a*∈ *{x, y, z }*. For a target miscoverage level *α*∈ (0, 1), We seek joint coverage while minimizing the prediction-box volume:

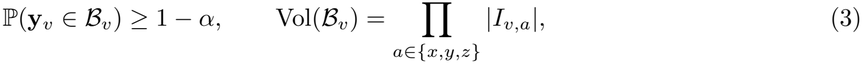

where |*I*_*v,a*_| is the interval length. Joint coverage requires all three coordinates to be covered simultaneously.

## 4 Methodology

We propose a dependence-aware CP framework for constructing 3D atom-level uncertainty regions in molecular docking (Figure 2). The framework addresses three challenges: heterogeneous prediction errors across molecular environments, statistical dependence among spatial coordinates, and distribution shifts between calibration and test data. We first estimate adaptive uncertainty intervals using a shared GNN, then calibrate these intervals jointly through copula-based dependence modeling. We further introduce DRE to adapt the calibration procedure under distribution shift.

To understand the challenges of uncertainty quantification in molecular docking, we analyze atomic prediction errors on the HiQBind 5K dataset (Wang et al., 2025). Figure 1 highlights two observations: (1) **Heterogeneous Prediction Errors** Prediction errors vary substantially across protein–ligand systems, indicating that uncertainty depends on both local atomic environments and global molecular context. This motivates our shared GNN for adaptive uncertainty estimation. (2) **Coordinate Dependence** Coordinate-wise nonconformity scores exhibit statistical dependence, suggesting that independent calibration may produce unnecessarily conservative 3D prediction regions. This motivates copula-based joint calibration. Additional error-distribution analyses are provided in Appendix L.

**Figure 1:**
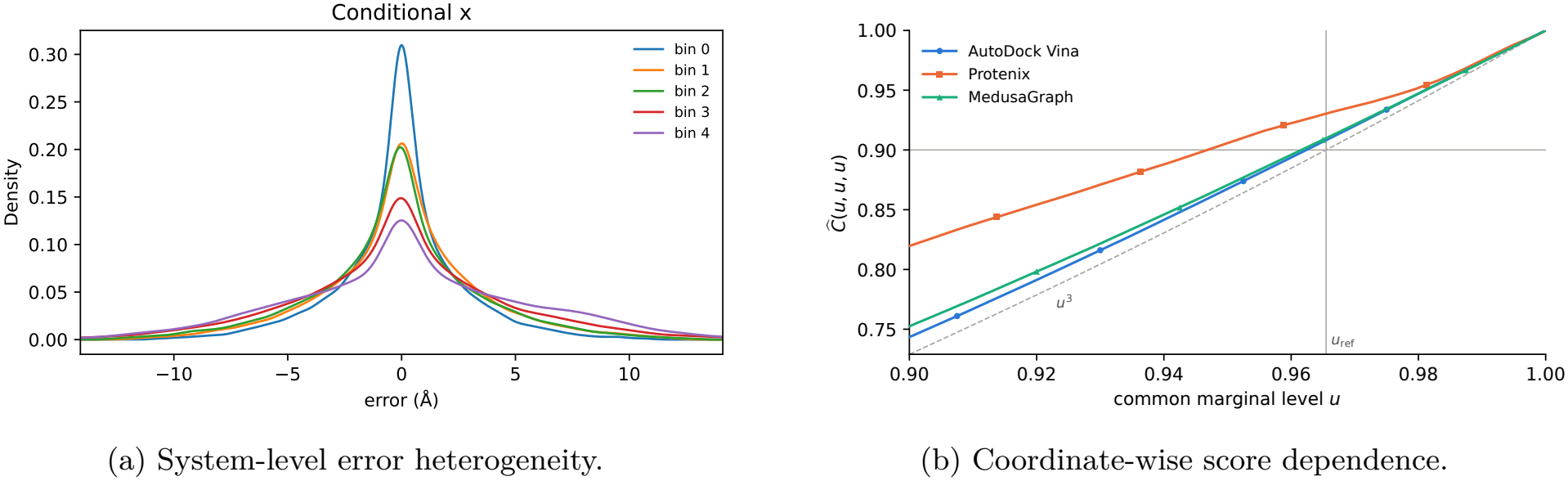
Motivating observations for dependence-aware uncertainty quantification. (a) Conditional signed *x*-coordinate error distributions for MedusaGraph, grouped into five bins by system-level mean prediction error, from lowest (bin 0) to highest (bin 4). (b) The empirical copula diagonal *Ĉ* (*u, u, u*) gives the probability that all three coordinate-wise nonconformity scores fall below their respective *u*-quantiles. Under independence this equals *u*^3^ (dashed). At *u*_ref_ = 0.90^1*/*3^ (vertical line) all three methods exceed the 90% joint coverage target (horizontal line), indicating conservative independence-based calibration due to dependence among coordinate-wise nonconformity scores.

**Figure 2:**
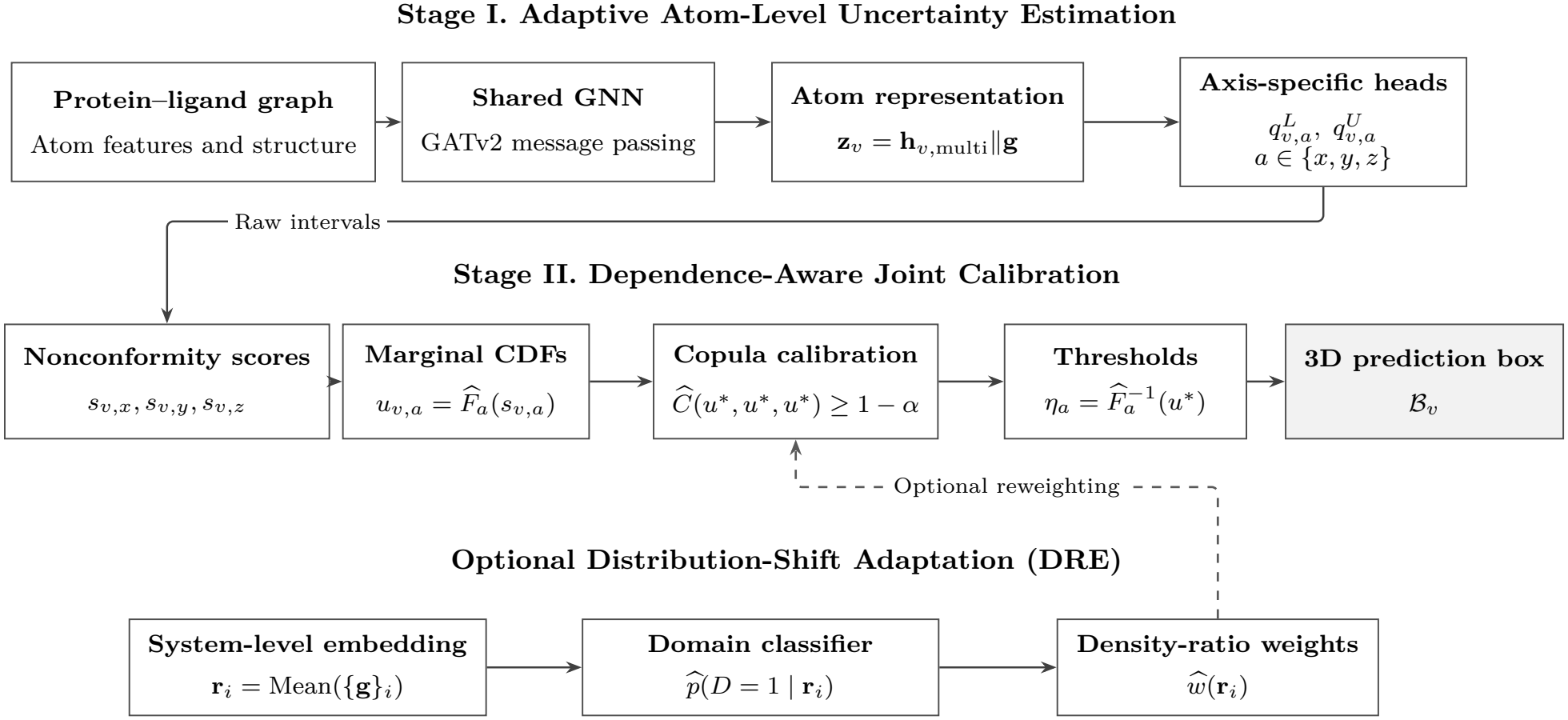
Overview of the proposed dependence-aware CP framework for molecular docking.

### 4.1 Shared GNN for Adaptive Uncertainty Estimation

Besides the heterogeneous prediction error shown in Figure 1a, local atomic features alone may not fully capture the difficulty of predicting an entire binding pose. We therefore develop a shared GNN that uses both local atomic environments and global molecular information to estimate atom-specific uncertainty.

#### Molecular Representation

Given a molecular graph, we use a shared GNN backbone with GATv2 message passing (Brody et al., 2022) to extract representations of individual atoms. For each atom *v*, the network processes its input feature vector **x**_*v*_ and aggregates information from neighboring atoms across multiple message-passing layers, producing a multiscale representation **h**_*v*,multi_. To capture broader molecular context, we obtain a graph-level representation **g** through global pooling and an encoder. The final representation of atom *v* is **z**_*v*_ = **h**_*v*,multi_ ∥**g**, where ∥denotes concatenation. This representation allows atoms with similar local structures to receive different uncertainty estimates depending on their molecular environments.

#### Asymmetric Uncertainty Estimation

Prediction errors need not be symmetric around the predicted coordinate. For example, molecular constraints may make displacement in one direction more likely than in the opposite direction. We therefore estimate separate lower and upper uncertainty widths for each spatial coordinate. Specifically, three axis-specific prediction heads share the molecular representation **z**_*v*_. For each axis *a* ∈ {*x, y, z*}, the corresponding head predicts a positive width scale *r*_*v,a*_ and an asymmetry parameter *ρ*_*v,a*_ ∈ (0, 1). The lower and upper widths are defined as

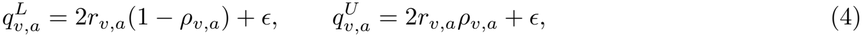

where *ϵ >* 0 ensures positive widths. The scale *r*_*v,a*_ controls interval size, while *ρ*_*v,a*_ determines how uncertainty is distributed between the lower and upper sides. These widths define the initial prediction interval

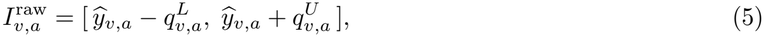

where *ŷ*_*v,a*_ is the coordinate predicted by the docking model.

We train the shared GNN and coordinate-specific heads using a quantile regression objective with width regularization. The quantile loss encourages intervals that reflect heterogeneous coordinate errors, while width regularization discourages unnecessarily large intervals. Importantly, this stage estimates uncertainty separately for each coordinate and does not enforce joint coverage. Network configurations, training objectives, and conformal-aware checkpoint selection are detailed in Appendix C.

### 4.2 Dependence-Aware Copula Calibration

Although the shared GNN provides adaptive uncertainty intervals, independently estimating uncertainty along the three coordinate axes does not ensure that the resulting 3D box contains the true atomic position. For example, even if each coordinate interval achieves 90% coverage, their simultaneous coverage may be substantially lower. A conservative alternative is to calibrate each coordinate at a higher coverage level, but this can produce unnecessarily large prediction boxes. The **key observation** is that coordinate errors are not necessarily independent (Figure 1b). A displacement caused by an incorrect ligand orientation, for instance, may simultaneously affect multiple coordinates. Modeling this dependence can therefore help determine how much calibration is needed to achieve a target joint coverage level without excessive conservatism.

#### Coordinate-Wise Nonconformity Scores

We first measure how well the raw uncertainty intervals cover the observed coordinates. For each calibration atom *v* and coordinate axis *a*, let *L*_*v,a*_ and *U*_*v,a*_ denote the endpoints of its raw prediction interval. We define

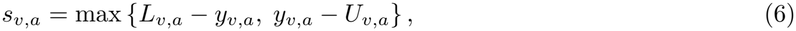

where *y*_*v,a*_ is the ground-truth coordinate. A positive score indicates that the true coordinate falls outside the interval, while a nonpositive score indicates that it is covered. Larger scores therefore indicate greater calibration corrections are needed. Because score distributions may differ across axes, we transform them onto a common probability scale. Let 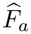 denote the empirical CDF of calibration scores for axis *a*. The transformed score is 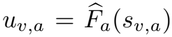, which indicates the relative position of a score within its marginal distribution. This transformation allows us to examine dependence among coordinates separately from differences in their score scales.

#### Joint Calibration Using Copulas

A copula models the joint dependence structure among the transformed coordinate scores. We investigate three copula models for dependence-aware calibration: Empirical, Gaussian, and R-vine copulas, which differ in their flexibility and assumptions about dependence. The Empirical Copula estimates dependence directly from calibration observations, while the Gaussian Copula models dependence through a correlation matrix of transformed scores, and the R-vine Copula captures more flexible dependence structures using pair-copula constructions.

Let *Ĉ* denote the estimated copula. Rather than selecting three independent calibration levels, we seek a common probability threshold *u*^*\**^ such that

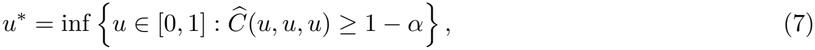

where 1− *α* is the target joint coverage level. Intuitively, *Ĉ* (*u, u, u*) estimates the probability that all three transformed scores fall below *u*. The threshold *u*^*\**^ therefore determines the marginal calibration levels needed to reach the desired joint coverage according to the fitted dependence model.

We transform this probability threshold back to each coordinate’s original score scale using 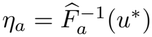, where 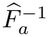 denotes the empirical quantile function. The final prediction box is then

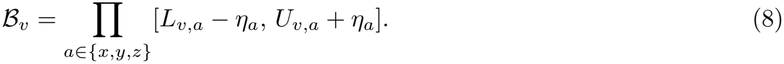

Here, *η*_*a*_ represents the calibration correction for coordinate *a*. By accounting for joint score dependence, the method aims to achieve the target joint coverage with smaller prediction boxes than conservative coordinate-wise calibration. For each copula model, we estimate the joint calibration level *u*^*\**^ using Eq. (7) and obtain axis-specific thresholds *η*_*a*_. Detailed formulations are provided in Appendix E.

Our implementation targets empirical joint coverage but does not directly establish a standard finite-sample test-coverage guarantee. We discuss coverage validity in Appendix H.

### 4.3 Distribution-Shift Adaptive Calibration

The preceding copula-based calibration relies on calibration scores being representative of test scores. However, molecular docking models may encounter proteins, ligands, or binding environments that differ from those in the calibration data, potentially altering the distribution of nonconformity scores and compromising joint coverage. To address this challenge, we introduce DRE to assign greater importance to calibration samples that are more representative of the test distribution.

#### DRE

Following classifier-based density ratio estimation (Bickel et al., 2009; Sugiyama et al., 2012), We represent each protein–ligand system *i* using a system-level embedding **r**_*i*_, obtained by averaging the graph-level representations **g** across its docking poses. A binary domain classifier distinguishes calibration systems (*D* = 0) from test systems (*D* = 1) based on these representations. The estimated density ratio is

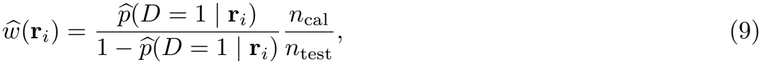

where 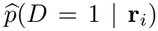 is the predicted test-domain probability, and *n*_cal_ and *n*_test_ denote the numbers of calibration and test systems used for domain classification. Larger weights indicate that a calibration system is more representative of the test distribution relative to the calibration distribution. Importantly, this procedure requires only unlabeled test representations, without accessing test coordinates or prediction errors.

#### Importance-Weighted Joint Calibration

Unlike standard weighted CP (Tibshirani et al., 2019), which adjusts calibration using importance weights, our approach introduces the estimated weights specifically at the copula calibration stage while retaining the original unweighted marginal score distributions. This design adjusts the joint calibration threshold based on the test-domain representation without modifying the marginal score transformations. Using the empirical copula as an example, the weighted copula diagonal is

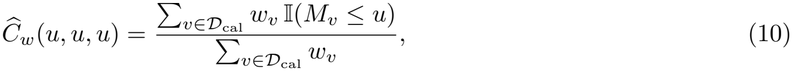

where *D*_cal_ denotes the set of calibration atoms, 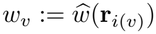 is the density ratio Eq. (9) of the system *i*(*v*) that contains atom *v, M*_*v*_ = max_*a*_ *u*_*v,a*_ is the maximum transformed score of Section 4.2, and I(·) is the indicator function. We determine the weighted joint calibration threshold as 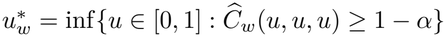, the analogue of Eq. (7), and construct the final prediction box via Eq. (8) with 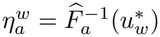 in place of *η*_*a*_. This procedure gives greater influence to calibration atoms from systems that are more representative of the test distribution. For Gaussian and R-vine copulas, the weights are incorporated into correlation estimation and likelihood-based fitting, respectively. To avoid unnecessary reweighting when distribution differences are negligible, we employ a permutation-based shift-detection procedure and retain uniform weights when no significant shift is detected. Details on classifier selection, shift detection, and weight stabilization are provided in Appendix F. The adaptation may improve empirical joint coverage under distribution shift but offers no general coverage guarantee (Tibshirani et al., 2019; Barber et al., 2023).

## 5 Experimental Setup

### 5.1 Dataset and Data Splits

Following Hao et al. (2026), we conduct experiments on the HiQBind 5K dataset (Wang et al., 2025), which contains protein–ligand complexes and docking poses generated by three representative molecular docking methods: AutoDock Vina, Protenix, and MedusaGraph. Each protein–ligand pair is treated as a system, with its docking poses represented as molecular graphs. HiQBind 5K enables consistent evaluation across multiple docking methods using the same underlying molecular systems. We further consider random and molecular-group splits, as well as controlled distribution shifts to investigate prediction-region efficiency and calibration reliability under different evaluation conditions.

#### Random Split

We select 4,496 protein–ligand systems shared by all three docking methods. Using a fixed random seed of 42, we partition these systems into training, validation, calibration, and test sets containing 3,147, 450, 450, and 449 systems, respectively. Splitting is performed at the system level, ensuring that all poses from the same system belong to the same subset to prevent information leakage.

#### Molecular Group Split

To evaluate generalization across distinct molecular systems, we construct a group-based split using PLINDER (Durairaj et al., 2024), a resource for organizing protein–ligand complexes by biological similarity. Of the 4,496 systems, 2,694 can be mapped to PLINDER. We group systems that share related protein identities, structural identifiers, or ligand structures, identified using SMILES (Simplified Molecular Input Line Entry System), a text-based representation of molecular structures (Weininger, 1988). Groups are defined transitively, ensuring that related systems remain in the same subset and preventing overlap across training, validation, calibration, and test sets. The resulting 217 groups are partitioned by assigning the seven largest to training, the next ten to validation, and the remaining 200 equally to calibration and test using seed 42. Because groups can contain different protein families, we refer to them as molecular groups rather than protein families. Although calibration and test groups are partitioned symmetrically, differences in their molecular compositions may still induce distribution shifts.

#### Controlled Split

To explicitly evaluate calibration under distribution shift, we construct controlled shifts by splitting 899 held-out systems to calibration and test sets based on system-level covariates. The trained predictors remain fixed, and the split mechanism varies the direction and strength of the shift.

### 5.2 Molecular Docking Backbones and Baseline

#### Molecular Docking Backbones

We evaluate three molecular docking methods representing distinct modeling paradigms. *AutoDock Vina* (Trott & Olson, 2010) uses conformational search within a predefined binding pocket and an empirical scoring function to rank ligand poses. *MedusaGraph* (Jiang et al., 2022) uses a graph neural network to predict ligand coordinates from an initial pose within the binding pocket. Protenix (ByteDance AML AI4Science Team et al., 2025) adopts an AlphaFold3-style diffusion-based framework for protein–ligand complex structure prediction. Following Hao et al. (2026), Vina and MedusaGraph operate within specified binding pockets, whereas Protenix must also infer the binding site, which may contribute to differences in their error distributions.

#### Baseline and Metrics

We compare our dependence-aware calibration methods—Empirical Copula, Gaussian Copula, and R-vine Copula against three baselines: *ConfDock, ConfDock (Bonferroni)* and *Independent. ConfDock* (Hao et al., 2026) uses a GNN to estimate asymmetric uncertainty widths for individual atomic coordinates, followed by coordinate-wise conformal calibration. Our methods are built upon ConfDock. *ConfDock (Bonferroni)* uses the same predictor but replaces this reference level with *u*_Bonf_ = 1 *−α/*3 *≈*0.96667. By the union bound, Bonferroni correction does not require independence between coordinate-wise coverage events, provided the marginal coverage requirements hold. *Independent* uses our shared GNN predictor but calibrates the three coordinates independently. For both baselines, we set the marginal calibration level to *u*_ref_ = 0.90^1*/*3^ *≈* 0.96549, corresponding to a 90% joint coverage target under the assumption of independent coordinate errors. All methods retain a 90% coordinate-wise training target, with the joint coverage adjustment applied during calibration. We evaluate their effectiveness in terms of achieving empirical joint coverage and prediction efficiency, i.e., prediction-box volume (the smaller, the better).

## 6 Results

### 6.1 Results for Random Split

Table 2 shows the results over 100 calibration/test re-divisions with the trained models fixed. Both ConfDock and Independent achieve average test joint coverage above the nominal 90% level, particularly for Protenix, where coverage reaches 92.85% and 93.31%, respectively. The overcoverage suggests that both baselines are overly conservative, resulting in unnecessarily large prediction boxes.

**Table 1:** Numbers of protein–ligand systems in the random and Molecular Group Splits.

| **Split** | **Train** | **Val** | **Calib** | **Test** | **Total** |
|---|---|---|---|---|---|
| Random | 3147 | 450 | 450 | 449 | 4496 |
| Molecular Group | 1998 | 206 | 247 | 243 | 2694 |

**Table 2:**
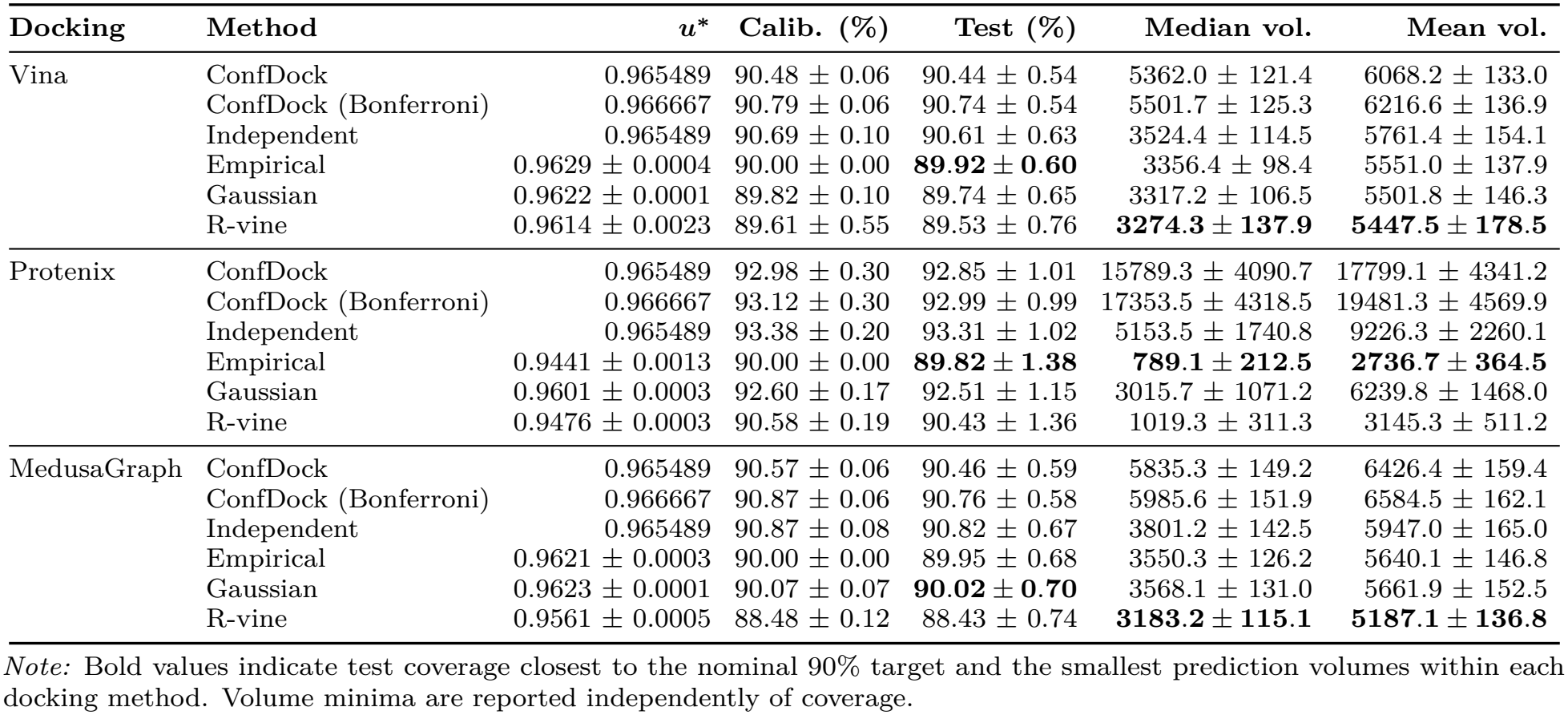
Random-split results. Entries are mean± standard deviation over 100 repetitions. Volumes are in Å^3^. Best results are in bold.

| **Docking** | **Method** | ***u*<sup>\*</sup>** | **Calib. (%)** | **Test (%)** | **Median vol.** | **Mean vol.** |
|---|---|---|---|---|---|---|
| Vina | ConfDock | 0.965489 | 90.48 ± 0.06 | 90.44 ± 0.54 | 5362.0 ± 121.4 | 6068.2 ± 133.0 |
|  | ConfDock (Bonferroni) | 0.966667 | 90.79 ± 0.06 | 90.74 ± 0.54 | 5501.7 ± 125.3 | 6216.6 ± 136.9 |
|  | Independent | 0.965489 | 90.69 ± 0.10 | 90.61 ± 0.63 | 3524.4 ± 114.5 | 5761.4 ± 154.1 |
|  | Empirical | 0.9629 ± 0.0004 | 90.00 ± 0.00 | **89.92 ± 0.60** | 3356.4 ± 98.4 | 5551.0 ± 137.9 |
|  | Gaussian | 0.9622 ± 0.0001 | 89.82 ± 0.10 | 89.74 ± 0.65 | 3317.2 ± 106.5 | 5501.8 ± 146.3 |
|  | R-vine | 0.9614 ± 0.0023 | 89.61 ± 0.55 | 89.53 ± 0.76 | **3274.3 ± 137.9** | **5447.5 ± 178.5** |
| Protenix | ConfDock | 0.965489 | 92.98 ± 0.30 | 92.85 ± 1.01 | 15789.3 ± 4090.7 | 17799.1 ± 4341.2 |
|  | ConfDock (Bonferroni) | 0.966667 | 93.12 ± 0.30 | 92.99 ± 0.99 | 17353.5 ± 4318.5 | 19481.3 ± 4569.9 |
|  | Independent | 0.965489 | 93.38 ± 0.20 | 93.31 ± 1.02 | 5153.5 ± 1740.8 | 9226.3 ± 2260.1 |
|  | Empirical | 0.9441 ± 0.0013 | 90.00 ± 0.00 | **89.82 ± 1.38** | **789.1 ± 212.5** | **2736.7 ± 364.5** |
|  | Gaussian | 0.9601 ± 0.0003 | 92.60 ± 0.17 | 92.51 ± 1.15 | 3015.7 ± 1071.2 | 6239.8 ± 1468.0 |
|  | R-vine | 0.9476 ± 0.0003 | 90.58 ± 0.19 | 90.43 ± 1.36 | 1019.3 ± 311.3 | 3145.3 ± 511.2 |
| MedusaGraph | ConfDock | 0.965489 | 90.57 ± 0.06 | 90.46 ± 0.59 | 5835.3 ± 149.2 | 6426.4 ± 159.4 |
|  | ConfDock (Bonferroni) | 0.966667 | 90.87 ± 0.06 | 90.76 ± 0.58 | 5985.6 ± 151.9 | 6584.5 ± 162.1 |
|  | Independent | 0.965489 | 90.87 ± 0.08 | 90.82 ± 0.67 | 3801.2 ± 142.5 | 5947.0 ± 165.0 |
|  | Empirical | 0.9621 ± 0.0003 | 90.00 ± 0.00 | 89.95 ± 0.68 | 3550.3 ± 126.2 | 5640.1 ± 146.8 |
|  | Gaussian | 0.9623 ± 0.0001 | 90.07 ± 0.07 | **90.02 ± 0.70** | 3568.1 ± 131.0 | 5661.9 ± 152.5 |
|  | R-vine | 0.9561 ± 0.0005 | 88.48 ± 0.12 | 88.43 ± 0.74 | **3183.2 ± 115.1** | **5187.1 ± 136.8** |

*Note:* Bold values indicate test coverage closest to the nominal 90% target and the smallest prediction volumes within each docking method. Volume minima are reported independently of coverage.

Among the three copula methods, Empirical Copula achieves the most consistent balance between coverage and prediction efficiency. Its test joint coverage is 89.92 ±0.60%, 89.82 ±1.38%, and 89.95± 0.68% for Vina, Protenix, and MedusaGraph, respectively, remaining close to the 90% target. Compared with ConfDock, it reduces median prediction-box volume by 37.40%, 95.00%, and 39.16%, respectively. These reductions arise from both the shared GNN predictor and dependence-aware calibration, as reflected by the additional improvements over the Independent baseline. ConfDock (Bonferroni) uses the more conservative level 1 −*α/*3, increasing test coverage to 90.74%, 92.99%, and 90.76% for Vina, Protenix, and MedusaGraph, respectively, at the cost of larger prediction boxes.

The Gaussian and R-vine copulas exhibit different coverage–efficiency trade-offs. Gaussian Copula performs comparably to Empirical Copula for Vina and MedusaGraph but remains conservative for Protenix, achieving 92.51% test coverage and substantially larger prediction boxes. R-vine Copula produces smaller boxes than Empirical Copula for Vina and MedusaGraph, but its coverage falls to 89.53% and 88.43%, respectively. For Protenix, R-vine achieves 90.43% coverage but produces larger boxes than Empirical Copula.

#### Dependence Analysis

To understand the differences among copula methods, we examine dependence between coordinate-wise nonconformity scores. Table 3 reports pairwise Kendall’s *τ*, while Figure 1b compares the diagonal sections of the fitted copulas. For coordinate axes *a, b* ∈ {*x, y, z*} with *a* = *b*, Kendall’s *τ* measures the concordance between their nonconformity scores: Positive values indicate that larger nonconformity scores along one axis tend to occur with larger scores along the other.

**Table 3:** Dependence among coordinate-wise nonconformity scores on the original random-split calibration set. Brackets denote 95% system-level bootstrap intervals from 200 resamples.

| Docking | τ<sub>xy</sub> | τ<sub>xz</sub> | τ<sub>yz</sub> | Mean τ | $\widehat{C}(u_{\mathrm{ref}})$ (%) |
|---|---|---|---|---|---|
| Vina | 0.25 | 0.23 | 0.24 | 0.24 | 90.82 |
|  |  |  |  | [0.22, 0.26] | [90.50, 91.14] |
| Protenix | 0.32 | 0.32 | 0.32 | 0.32 | 93.05 |
|  |  |  |  | [0.29, 0.34] | [92.44, 93.55] |
| MedusaGraph | 0.25 | 0.22 | 0.24 | 0.24 | 90.98 |
|  |  |  |  | [0.22, 0.26] | [90.69, 91.25] |

All three docking methods exhibit positive coordinate dependence, with mean pairwise Kendall’s *τ* ranging from 0.24 to 0.32. Protenix shows the strongest dependence (*τ* = 0.32), compared with Vina and MedusaGraph (both approximately 0.24). This helps explain why independence-based calibration is particularly conservative for Protenix. We further examine the empirical joint coverage at the independence-based marginal threshold *u*_ref_ = 0.90^1*/*3^. As shown in Table 3, the coverage of the empirical copula diagonal for all three docking methods exceeds the nominal 90% target. The larger deviation for Protenix is consistent with its stronger coordinate dependence and the substantial volume reductions achieved by dependence-aware calibration. These findings demonstrate that accounting for coordinate dependence can reduce conservative calibration. They also highlight the importance of estimating the joint score distribution accurately: more flexible copula models do not necessarily provide better coverage–efficiency trade-offs.

#### Coverage–Volume Analysis

To disentangle the effects of uncertainty estimation and joint calibration, we examine prediction-box volumes across different calibration thresholds while keeping the trained predictors and test systems fixed. Figure 3 presents the resulting coverage–volume trade-offs for the three docking methods. At matched empirical test coverage levels between 90% and 94%, the shared GNN consistently produces smaller median prediction-box volumes than ConfDock. At 90% test coverage, the volume reductions are 35.54%, 70.42%, and 36.66% for Vina, Protenix, and MedusaGraph, respectively. These post-hoc comparisons indicate that the shared GNN provides more efficient uncertainty estimates independently of the calibration strategy. The Independent baseline and all three unweighted copula methods follow the same coverage–volume curve because they share the same predictor and marginal score transformations. Their distinction lies in selecting different calibration thresholds *u*^*\**^, which determine the operating points along this curve. Empirical Copula accounts for coordinate dependence to select thresholds that achieve coverage closer to the nominal 90% target without unnecessary conservatism. ConfDock (Bonferroni) uses a more conservative calibration threshold, achieving slightly higher joint coverage at the cost of larger prediction-box volumes. This shifts the operating point along the original ConfDock curve without improving the underlying uncertainty estimator.

**Figure 3:**
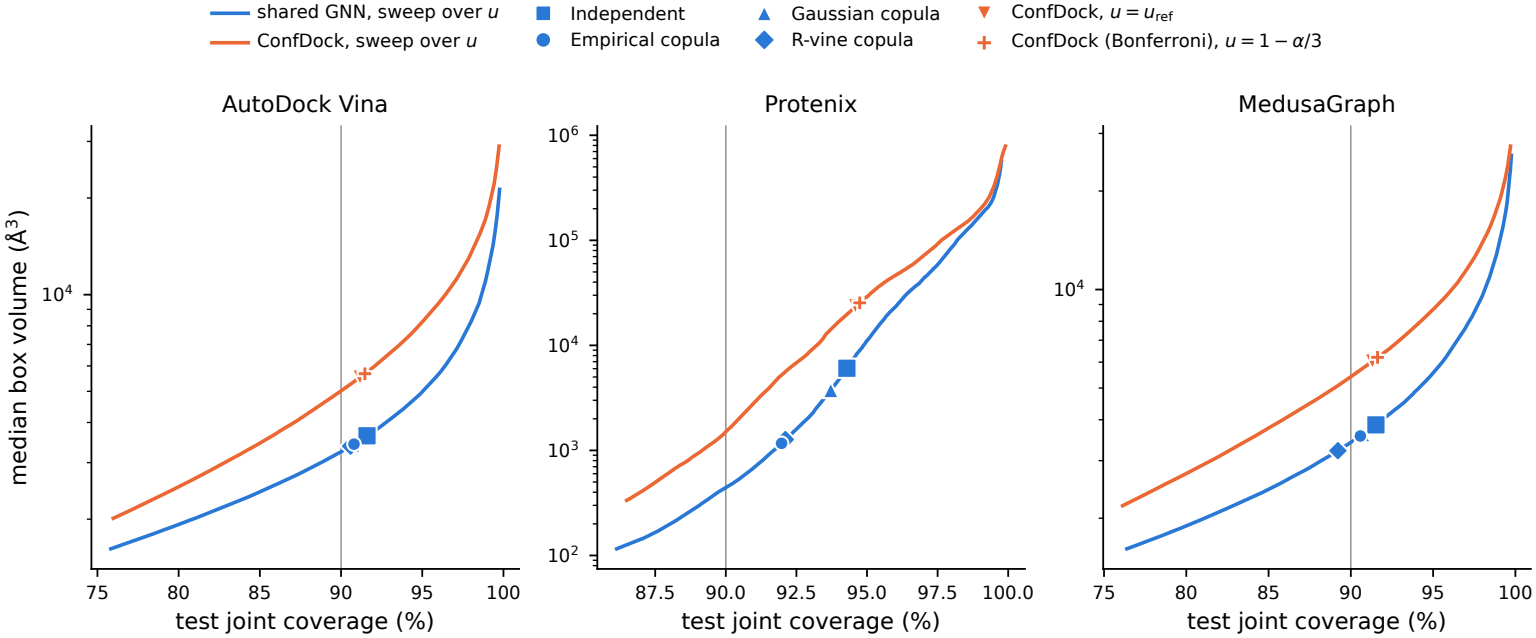
Coverage–volume trade-offs for three docking methods on the original split. Blue and orange curves correspond to the shared GNN and ConfDock predictors. Markers indicate calibration operating points, and the vertical line denotes 90% test joint coverage. Coverage-matched comparisons are post-hoc diagnostics.

This effect is particularly pronounced for Protenix, where median prediction-box volume increases from approximately 449 Å^3^ to 4,877 Å^3^ as test coverage rises from 90% to 94%. This steep increase is consistent with Protenix’s bimodal docking-error distribution, whose broad high-error component may cause calibration quantiles to increase rapidly at higher coverage levels. Since prediction-box volume depends multiplicatively on the three coordinate-wise interval widths, larger calibration corrections can substantially amplify the volume increase (Appendix J.1). Consequently, small differences in calibration thresholds can produce substantial changes in prediction-box volume. Gaussian Copula remains relatively conservative for Protenix, while R-vine sometimes produces smaller boxes at the expense of undercoverage, as shown in Table 2. Overall, these results reveal two complementary contributions: the shared GNN improves the coverage–volume trade-off, while dependence-aware calibration selects more efficient operating points. Additional calibration sensitivity results are provided in Appendix Figure 4.

### 6.2 Results under Controlled Distribution Shifts

To evaluate the effectiveness of DRE under distribution shift, we construct controlled covariate shifts while keeping the trained docking predictor fixed. We focus on Protenix, which exhibits the largest coverage-relevant shifts in a preliminary validation analysis. We consider six shift directions, including three individual and three combined covariate shifts, with shift strength controlled by *β*, where its sign determines the direction and its magnitude determines the strength of the shift. We compare standard unweighted calibration (Standard), calibration using known density ratios (Oracle), and calibration using estimated density ratios (DRE). Each setting is repeated 30 times with a target joint coverage of 90%. Details of the shift construction and validation-based screening are provided in Appendix G.

#### Coverage under Distribution Shift

Table 4 reports the signed test coverage gap relative to the 90% target, where negative values indicate undercoverage and positive values indicate overcoverage. Across 36 controlled-shift settings, DRE reduces the average absolute coverage gap from 3.01 to 0.80 percentage points (pp), a 73.4% reduction. Among the 21 settings where Standard (Std.) exhibits an absolute mean coverage gap exceeding 2 pp, DRE improves all 21.

**Table 4:** Coverage gaps (pp) and mean median prediction volumes (Å^3^) under individual and combined controlled shifts for Protenix.

| **Panel A: Individual shift directions** |  |  |  |  |  |  |  |  |  |
|---|---|---|---|---|---|---|---|---|---|
|  | **PCA01** |  |  | **PCA19** |  |  | **PCA07** |  |  |
| *β* | Std. | Oracle | DRE | Std. | Oracle | DRE | Std. | Oracle | DRE |
| *Coverage gap (pp)* |  |  |  |  |  |  |  |  |  |
| −2.0 | −12.86 | −0.94 | −3.54 | −0.16 | −0.77 | +1.27 | +3.27 | −1.09 | −0.92 |
| −1.0 | −8.59 | −0.65 | −2.14 | −0.29 | +0.03 | +0.73 | +2.25 | −0.38 | +0.04 |
| −0.5 | −4.95 | −0.52 | −1.72 | −0.40 | −0.31 | −0.07 | +1.23 | −0.25 | +0.49 |
| +0.5 | +3.55 | −0.34 | +1.25 | +0.39 | +0.19 | +0.34 | −2.07 | −0.36 | −0.49 |
| +1.0 | +6.31 | +0.80 | +1.66 | +0.37 | −0.03 | +0.28 | −3.70 | −0.12 | +0.14 |
| +2.0 | +8.33 | +0.01 | +0.46 | −0.01 | −0.98 | +0.33 | −6.81 | −0.39 | −0.84 |
| *Mean median prediction volume (Å<sup>3</sup>)* |  |  |  |  |  |  |  |  |  |
| −2.0 | 243.3 | 19587.1 | 4558.7 | 750.0 | 4041.2 | 2072.5 | 1512.9 | 4123.5 | 258.4 |
| −1.0 | 285.1 | 6210.9 | 3925.7 | 770.5 | 1402.1 | 1417.7 | 1211.7 | 388.2 | 476.7 |
| −0.5 | 391.4 | 2187.2 | 1493.9 | 735.1 | 791.5 | 882.1 | 1026.9 | 527.2 | 822.6 |
| +0.5 | 1719.4 | 304.4 | 606.0 | 885.4 | 873.9 | 896.5 | 596.1 | 1084.5 | 1051.9 |
| +1.0 | 4522.0 | 241.0 | 314.3 | 867.1 | 914.5 | 1212.5 | 505.3 | 1669.3 | 2303.7 |
| +2.0 | 9760.4 | 186.9 | 126.1 | 815.3 | 13151.2 | 1181.2 | 369.4 | 5694.5 | 3055.7 |
| **Panel B: Combined shift directions** |  |  |  |  |  |  |  |  |  |
|  | **Linear** |  |  | **Interaction** |  |  | **Nonlinear** |  |  |
| *β* | Std. | Oracle | DRE | Std. | Oracle | DRE | Std. | Oracle | DRE |
| *Coverage gap (pp)* |  |  |  |  |  |  |  |  |  |
| −2.0 | −2.79 | −1.93 | +0.47 | −0.41 | −0.48 | +0.62 | +6.17 | −0.21 | +1.09 |
| −1.0 | −2.16 | −0.46 | +1.01 | +0.09 | −0.22 | +0.59 | +4.39 | −0.10 | +1.39 |
| −0.5 | −1.46 | −0.14 | −0.01 | +0.06 | −0.36 | +0.07 | +2.88 | +0.34 | +1.56 |
| +0.5 | +1.06 | −0.39 | +0.13 | −0.41 | −0.18 | −0.07 | −3.06 | −0.37 | −1.33 |
| +1.0 | +2.11 | −0.04 | −0.08 | −0.07 | +0.20 | +0.07 | −5.07 | +0.07 | −1.40 |
| +2.0 | +2.87 | −1.09 | −0.52 | +0.24 | −1.40 | −0.37 | −7.65 | −0.38 | −1.34 |
| *Mean median prediction volume (Å<sup>3</sup>)* |  |  |  |  |  |  |  |  |  |
| −2.0 | 463.0 | 5813.0 | 4032.8 | 753.3 | 3428.7 | 1654.7 | 4111.2 | 237.2 | 277.5 |
| −1.0 | 517.3 | 1663.6 | 6981.5 | 817.3 | 842.3 | 1079.9 | 2259.9 | 284.7 | 495.9 |
| −0.5 | 613.4 | 1092.7 | 1737.1 | 819.1 | 719.3 | 836.5 | 1505.4 | 478.2 | 829.3 |
| +0.5 | 1029.3 | 589.1 | 712.8 | 756.7 | 865.6 | 904.0 | 488.1 | 1496.6 | 1137.0 |
| +1.0 | 1313.4 | 682.8 | 581.2 | 777.6 | 1009.6 | 945.3 | 392.5 | 4074.8 | 1688.6 |
| +2.0 | 1560.7 | 532.3 | 607.2 | 822.2 | 4738.9 | 775.9 | 291.0 | 22536.0 | 3516.7 |

The improvements are particularly pronounced when distribution shifts substantially affect coverage. The three individual shift directions, PCA01, PCA19, and PCA07, are derived from principal-component scores of system-level molecular representations. Their construction is detailed in Appendix G. For example, under PCA01 with *β* = −2, Standard coverage falls to 77.14%, while DRE increases it to 86.46%, approaching the Oracle coverage of 89.06%. Under the nonlinear shift with *β* = +2, DRE reduces the coverage gap from −7.65 to −1.34 pp. In contrast, PCA19 and Interaction shifts introduce relatively small coverage deviations, leaving limited room for improvement. These results indicate that DRE is most beneficial when covariate shifts induce substantial changes in the nonconformity-score distribution.

#### Coverage–Volume Trade-off

Improved calibration does not always produce smaller prediction regions. When Std. suffers from undercoverage, DRE generally enlarges prediction boxes (i.e., sacrifices prediction efficiency) to improve coverage. For example, under PCA01 with *β* = −2, the mean median volume increases from 243.3 to 4558.7 Å^3^. Conversely, when Std. overcovers, DRE can reduce both coverage deviation and prediction volume. Under PCA01 with *β* = +2, volume decreases from 9760.4 to 126.1 Å^3^. Thus, DRE primarily improves coverage reliability rather than uniformly reducing prediction volume.

#### Effect of Shift Strength

Stronger shifts lead to more distinguishable calibration and test distributions but also more variable importance weights, reducing the effective sample size. When no controlled shift is introduced (*β* = 0), the shift-detection gate generally avoids unnecessary reweighting, yielding nearly identical coverage to Standard calibration. Additional diagnostics are provided in Appendix G.3.

### 6.3 Results for Molecular Group Split

We further evaluate our method under the Molecular Group Split, which assesses generalization across distinct molecular groups. Although calibration and test systems are drawn from the same held-out pool, differences in their molecular compositions may lead to distribution shifts. Table 5 compares standard empirical-copula calibration across three docking methods and DRE-weighted calibration for Protenix. Under standard calibration, Protenix produces substantially larger prediction boxes than Vina and MedusaGraph, with a median volume approximately ten times greater. Several factors may contribute to this difference. First, Vina and MedusaGraph use predefined binding pockets, whereas Protenix also predicts the binding site, introducing an additional source of positional uncertainty. Protenix exhibits a heterogeneous error distribution, with mostly small errors but occasional large deviations. Such errors can increase the global conformal correction *η*_*a*_, enlarging prediction boxes even for atoms with relatively accurate predictions. Second, differences between training and calibration systems may limit the GNN’s ability to predict sufficiently large uncertainty widths for difficult, previously unseen molecular groups. Consequently, these errors may be addressed primarily through larger calibration corrections rather than adaptive atom-specific widths.

**Table 5:** Results for the Molecular Group Split with and without DRE. Prediction-box volumes are in Å^3^.

| **Docking** | **Calibration** | **Test (%)** | **Median vol.** | **Mean vol.** |
|---|---|---|---|---|
| MedusaGraph | Standard | 91.43 | 5563.11 | 7699.09 |
| Vina | Standard | 91.97 | 5373.70 | 7852.43 |
| Protenix | Standard | 92.31 | 57644.30 | 75977.14 |
| Protenix | DRE (*k* = 50) | **90.04** | **29271.04** | **42581.63** |

Third, the calibration and test sets differ in their composition of high-error systems, which may further affect calibration efficiency. MedusaGraph, Vina, and Protenix achieve test joint coverage of 91.43%, 91.97%, and 92.31%, respectively, indicating mild overcoverage. We focus the DRE analysis on Protenix, which exhibits the largest coverage deviation and prediction-box volumes. With DRE (*k* = 50), test coverage approaches the nominal target, decreasing from 92.31% to 90.04%, while median and mean prediction-box volumes decrease by 49.22% and 43.95%, respectively. These results suggest that DRE can improve calibration efficiency by adapting joint calibration thresholds to differences in the realized molecular compositions of calibration and test systems. Further analyses of docking-error distributions, training–calibration heterogeneity, calibration–test imbalance, and DRE sensitivity are provided in Appendix J.

### 6.4 Sensitivity to Weight Placement

Our distribution-shift adaptation applies density-ratio weights during copula-based joint calibration while retaining unweighted marginal score distributions (*frozen marginals*). An alternative is to also incorporate these weights into the marginal score distributions (*fully weighted*).

We compare both strategies using the empirical copula on Protenix across six controlled-shift scenarios, with *β* ∈{± 0.5, ±1, ± 2}. Each setting is repeated 30 times with a target joint coverage of 90%. We consider both oracle density ratios and estimated DRE weights; the two strategies use identical weights and differ only in whether the marginal score distributions are weighted. Detailed formulations and results are provided in Appendix K.

As shown in Table 6, under moderate shifts (|*β* |≤1), both strategies achieve similar coverage, although fully weighted calibration generally increases prediction-box volumes under positive moderate shifts without improving coverage. At *β* = +2, using oracle weights, fully weighted calibration reduces prediction-box volume but increases the absolute coverage deviation from 0.90 to 1.21 pp. In contrast, stabilized DRE weights yield more similar results across the two strategies. Since fully weighted marginals provide no consistent improvement in coverage or prediction efficiency, we maintain frozen marginals in our framework.

**Table 6:**
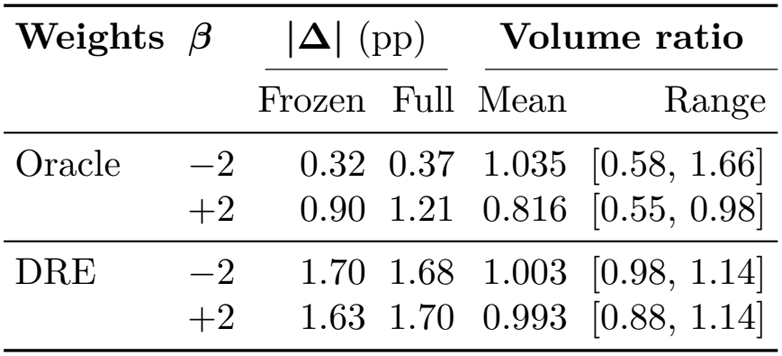
Weight placement under controlled shifts. |Δ| : coverage deviation (pp); volume ratio: full/frozen.

| **Weights** | ***β*** | \|**Δ**\| (pp) |  | **Volume ratio** |  |
|---|---|---|---|---|---|
|  |  | Frozen | Full | Mean | Range |
| Oracle | −2 | 0.32 | 0.37 | 1.035 | [0.58, 1.66] |
|  | +2 | 0.90 | 1.21 | 0.816 | [0.55, 0.98] |
| DRE | −2 | 1.70 | 1.68 | 1.003 | [0.98, 1.14] |
|  | +2 | 1.63 | 1.70 | 0.993 | [0.88, 1.14] |

## 7 Conclusions

We develop a dependence-aware CP framework for 3D atom-level uncertainty quantification in molecular docking, combining GNN-based adaptive uncertainty estimation, copula-based joint calibration, and DRE for distribution-shift adaptation.

Experiments across three docking methods demonstrate that dependence-aware calibration produces more compact prediction regions while maintaining empirical joint coverage near the target level. Empirical Copula achieves the most consistent balance between coverage reliability and prediction efficiency. Under the Molecular Group Split and controlled distribution shifts, DRE further improves calibration when distribution differences affect prediction errors, although its effectiveness depends on shift characteristics and estimation accuracy. Rare, large docking errors remain challenging, often leading to conservative calibration and enlarged prediction regions. Future work will explore conditional calibration methods for heterogeneous error regimes. Overall, our findings highlight the importance of accounting for coordinate dependence and distribution shifts in reliable molecular docking uncertainty quantification.

## Acknowledgments

This work is supported by the National Science Foundation (NSF) Grant #2621882, NSF #2621883, NSF ACCESS Computing Resources, NAIRR, and Argonne Leadership Computing Facility.

## Data Availability

The source code and analysis scripts used in this work are available at https://github.com/Guanyulyu/hybrid-v3-distribution-shift. The HiQBind 5K molecular docking graph datasets are available on Hugging Face at https://huggingface.co/datasets/liofoil/copuladock. Related ConfDock graph data resources are available at https://huggingface.co/datasets/liofoil/gnncp.

## Appendix

### A Algorithms

Algorithms 1 and 2 are the pseudocodes for Dependence aware GNN training part structure and the checkpoint strategy on the validation set.

#### Algorithm 1

Shared GNN Training for Coordinate-wise Uncertainty

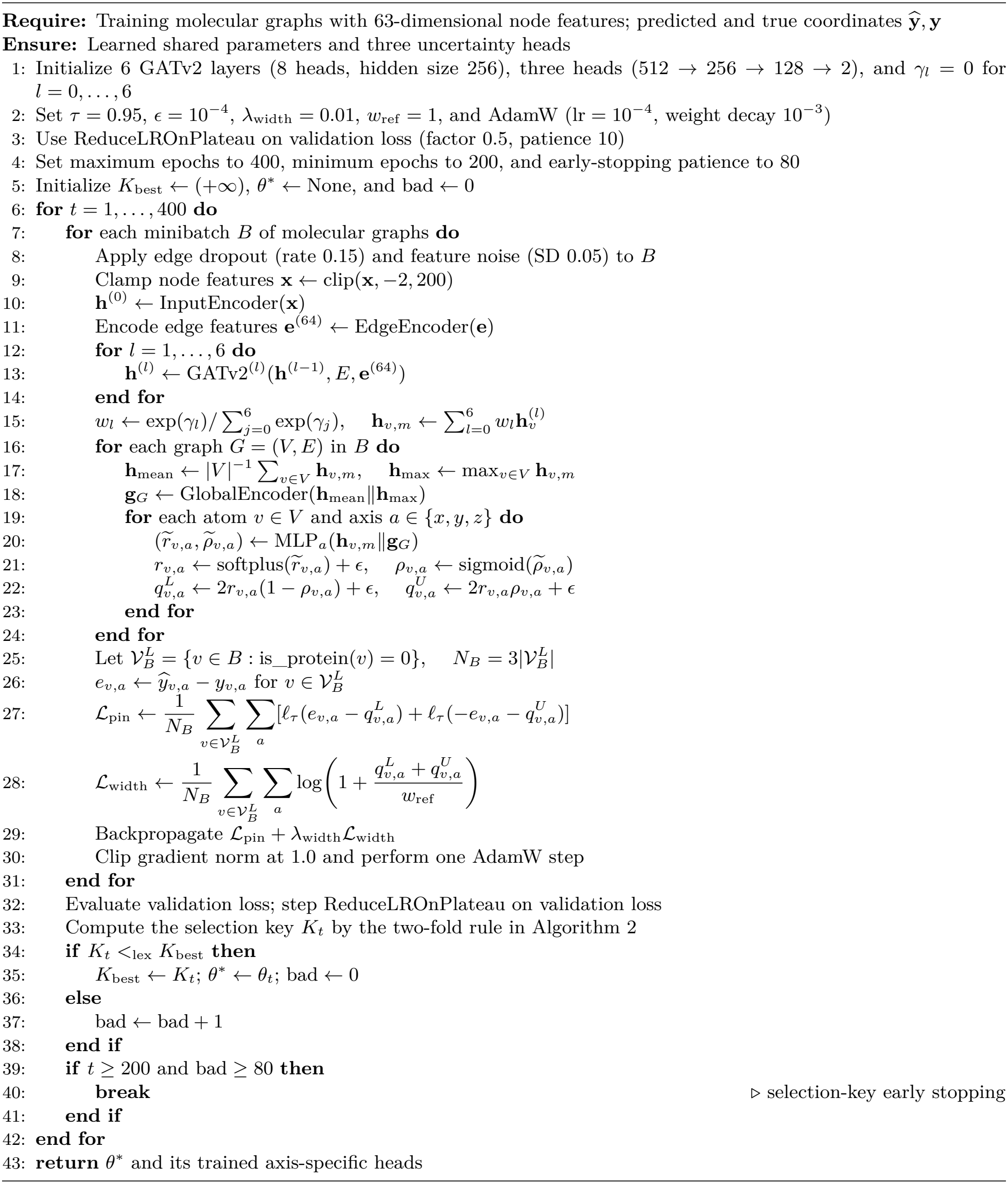

#### Algorithm 2

Two-Fold Conformal-Aware Checkpoint Selection

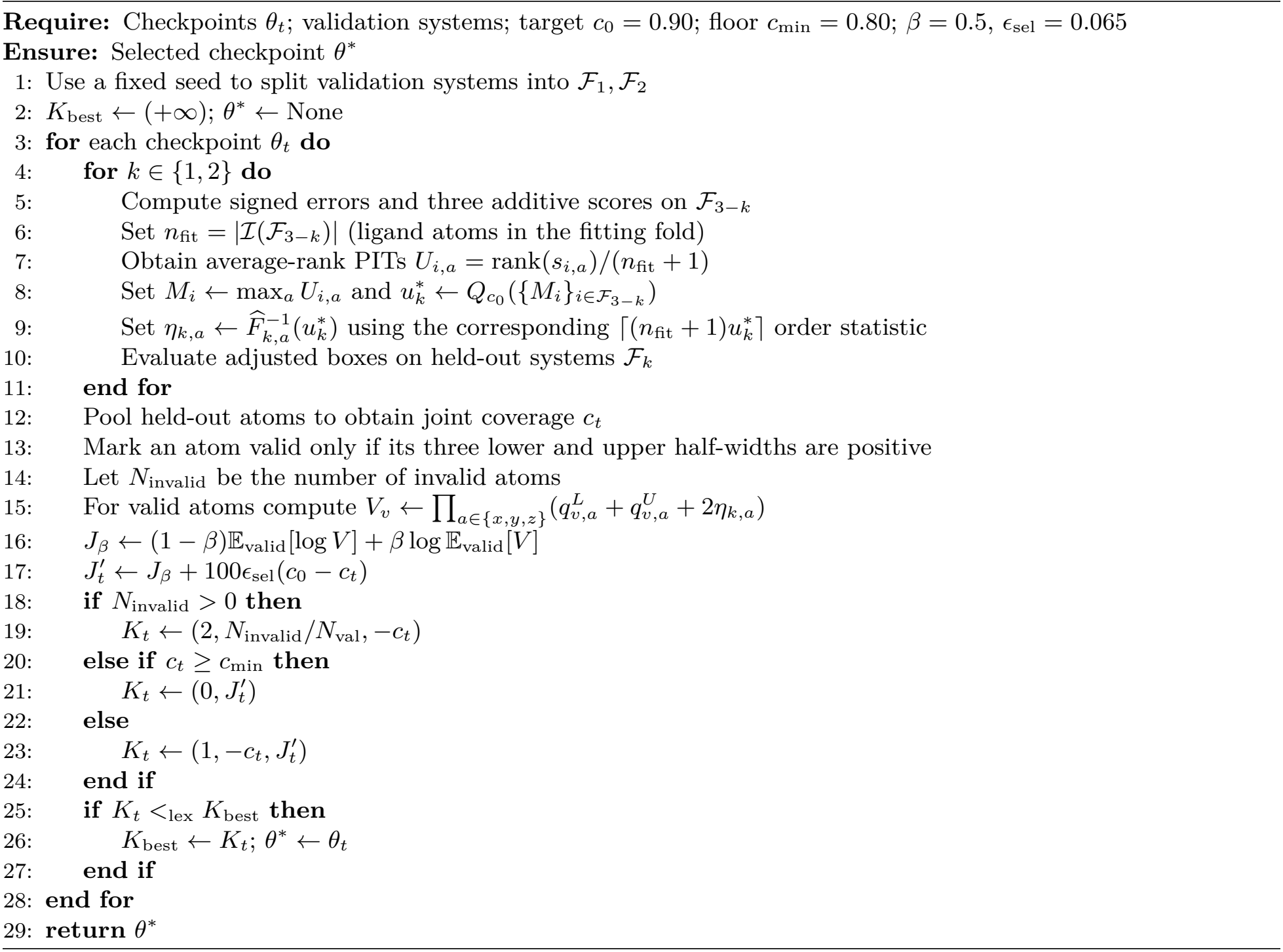

Algorithm 3 and Algorithm 4 are the pseudocodes for DRE and Copula Calibration.

#### Algorithm 3

Optional DRE: OOF Selection and Weight Stabilization

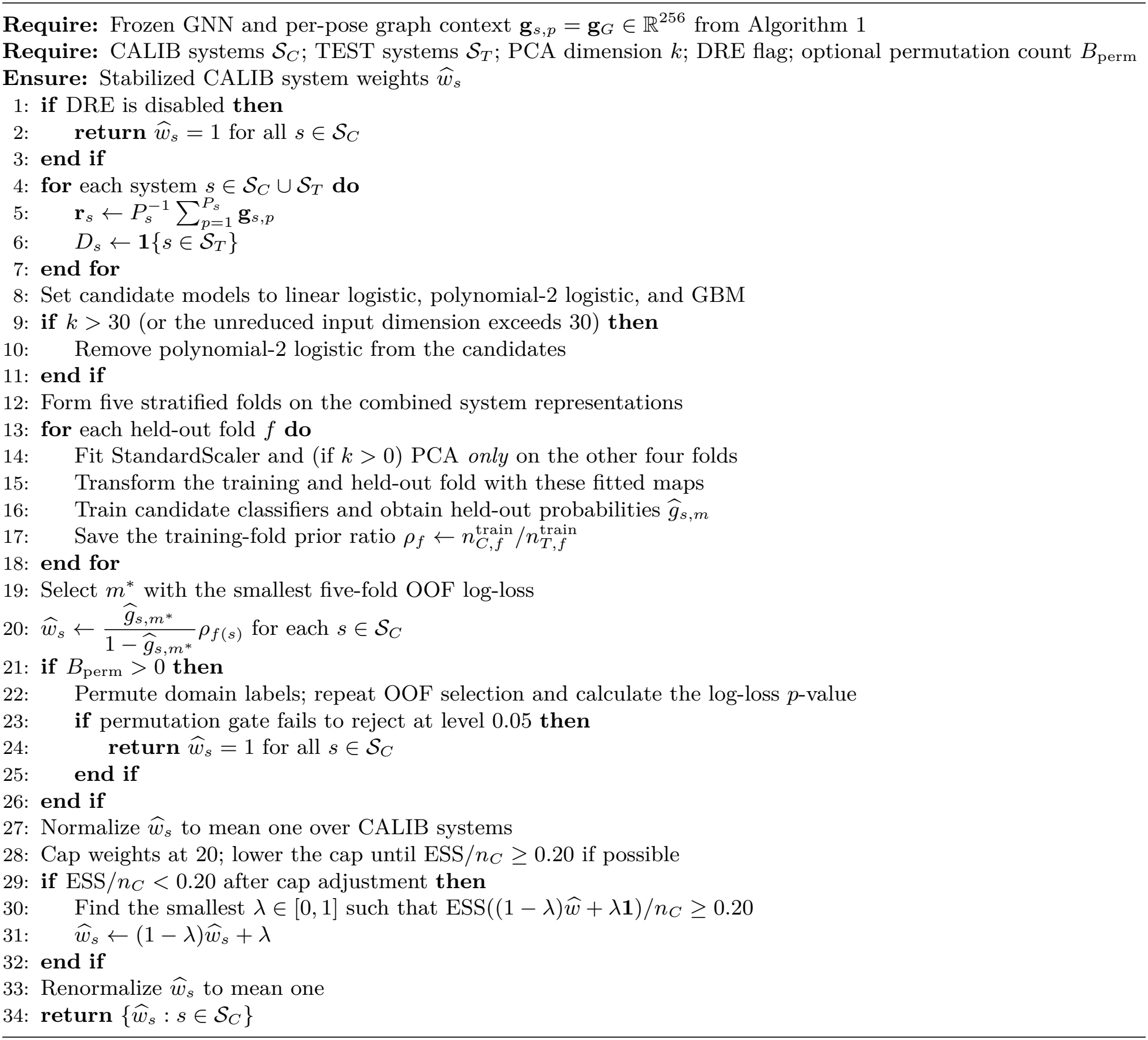

#### Algorithm 4

Copula Calibration with Frozen Unweighted Marginals

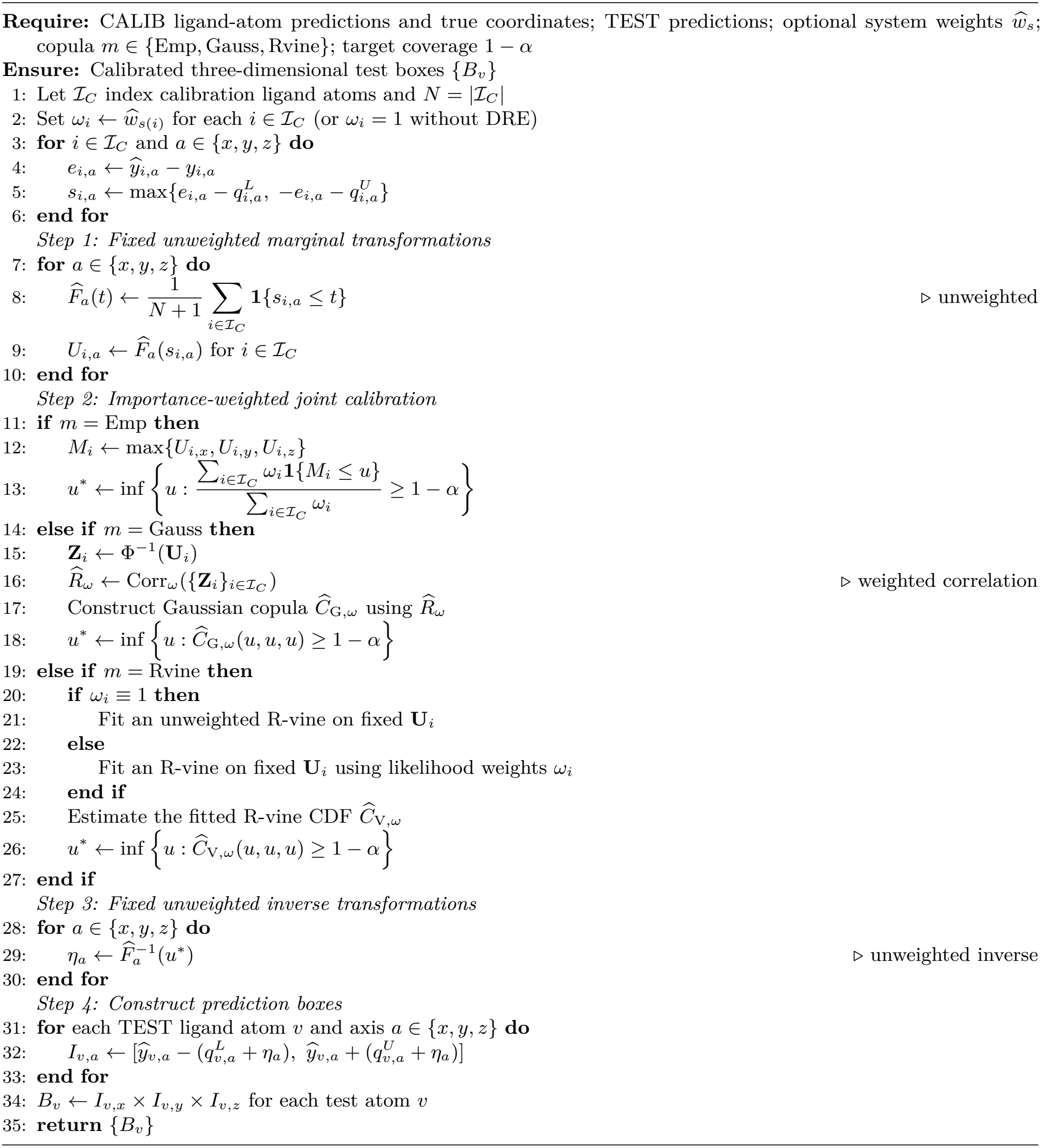

### B Additional Preliminaries

**Marginal and Joint Coverage** For each coordinate axis *a* ∈ {*x, y, z*}, marginal coverage requires

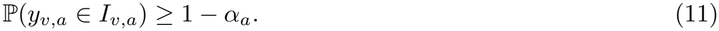

In contrast, joint coverage requires

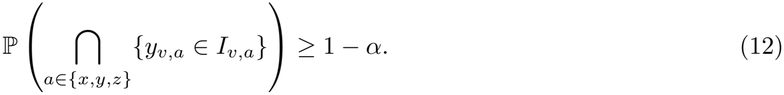

Marginal coordinate-wise coverage alone does not guarantee joint coverage at the same target level.

**Split Conformal Prediction** Split CP uses a trained prediction model and a held-out calibration set 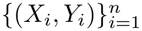 (Papadopoulos et al., 2002; Lei et al., 2018). A nonconformity score *s*(*X*_*i*_, *Y*_*i*_) measures the discrepancy between predictions and observations. Let 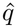 be the ⌈(*n* + 1)(1 − *α*)⌉-th smallest calibration score, with 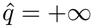 if the index exceeds *n*. The conformal prediction set for a new input *X*_*n*+1_ is

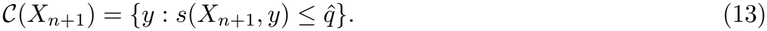

Under exchangeability,

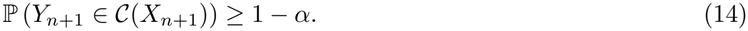

Absolute-residual scores yield symmetric intervals, whereas conformalized quantile regression supports adaptive and asymmetric intervals (Romano et al., 2019).

#### Exchangeability and Distribution Shift

Standard CP assumes exchangeability of calibration and test observations. For a scoring function fixed independently of these observations, this assumption implies exchangeability of the nonconformity scores, which underlies the finite-sample coverage guarantee (Vovk et al., 2005; Lei et al., 2018; Tibshirani et al., 2019). Exchangeability of observations is sufficient but not necessary for score exchangeability. Under covariate shift, where calibration and test distributions share *P*_*Y*_| _*X*_ but differ in *P*_*X*_, score distributions may remain identical if the conditional score distribution does not depend on *X*. More generally, coverage degradation depends on discrepancies between the calibration and test score distributions (Barber et al., 2023; Cauchois et al., 2024).

### C Shared Information GNN Architecture Details

#### C.1 Atom level Information

We use a high-dimensional (63 dims after selection) atom feature vector **x**_*v*_ as the input for each atom *v*. The input features contain both the information of the atom itself and information related to its local molecular environment.

Before the GATv2 message-passing layers, the input feature vector **x**_*v*_∈ ℝ^63^ is first transformed by an input encoder into a 256-dimensional hidden representation, The input encoder can be written as

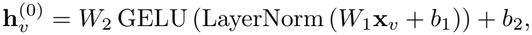

where **x**_*v*_ ∈ ℝ^63^ and 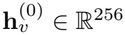.

The hidden representation 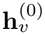 is then processed by six GATv2 layers with eight attention heads per layer, producing 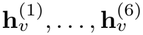. Since deeper layers may gradually lose some information from earlier stages, we keep the representations from different message-passing depths. The model uses learnable `scale_logits` and a softmax function to compute the importance of each layer. The multiscale representation is then written as

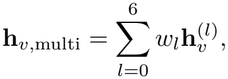

where the weights are obtained through a softmax function,

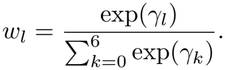

The learnable parameters *γ*_0_, …, *γ*_6_ are initialized to zero,

Therefore, at the beginning of training, the softmax gives equal weights to all seven representations,

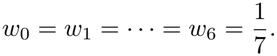

During training, the parameters *γ*_*l*_ are updated through backpropagation, allowing the model to learn the relative importance of information from different message-passing depths. Therefore, representations from both shallow and deep layers can contribute to the final shared feature, with their importance controlled by the learned weights.

##### C.1.1 Graph level Information

We then apply mean pooling and max pooling to the multiscale node representations **h**_*v*,multi_ to obtain graph-level information. For a molecular graph with node set *V*, the two pooled representations are

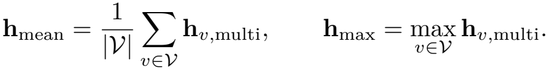

where the maximum is taken element-wise over all nodes. These two representations are concatenated and passed through a global encoder. Here **g** = *f*_global_ (**h**_mean_∥ **h**_max_), where ∥ denotes concatenation. The resulting graph-level representation **g** contains global information from the entire protein–ligand graph.

##### C.1.2 Information Combination

For each atom *v*, we have **z**_*v*_ that combine atom level information **h**_*v*,multi_ and graph level information **g**, with 512 dims as a result of 256 dims plus 256 dims. We have **z**_*v*_ = **h**_*v*,multi_∥**g**.

Therefore, the *x, y*, and *z* axes share the same molecular representation before reaching their axis-specific output heads.

#### C.2 The information that three axes does not share

Although the three coordinate axes share the same molecular representation, their uncertainty distributions can still be different. The prediction error along one axis does not necessarily have to be the same scale or asymmetry as the other two axes. Therefore, after the shared representation **z**_*v*_ is obtained, we use three independent output heads for the *x, y*, and *z* axes.

For each axis *a* ∈ {*x, y, z* }, we use an independent axis-specific output head. Each head is implemented as a small multilayer perceptron (MLP). It takes the shared representation **z**_*v*_ as input and maps it through hidden layers before producing two axis-specific quantities: a width scale 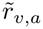 and an asymmetry parameter 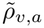

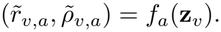

They are transformed as

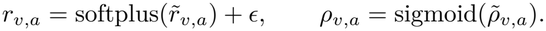

where *r*_*v,a*_ *>* 0 and *ρ*_*v,a*_ ∈ (0, 1).

The lower and upper raw uncertainty widths are then given by

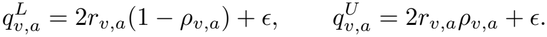

The combination of *r*_*v,a*_ and *ρ*_*v,a*_ provides two degrees of freedom for the predicted quantile.

Therefore, the three axes share the same molecular features, but each axis has its own output head to learn its uncertainty scale and asymmetry. No cross-axis constraint is applied on the raw widths at this stage.

#### C.3 Marginal Training Objective

After obtaining the lower and upper raw uncertainty widths 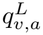 and 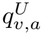, each axis is trained using a marginal quantile regression objective. For atom *v* and axis *a* ∈ {*x, y, z*}, we define the signed coordinate error as *e*_*v,a*_ = *ŷ*_*v,a*_ − *y*_*v,a*_.

where *ŷ*_*v,a*_ and *y*_*v,a*_ are the predicted and ground-truth coordinates, respectively.

For a target marginal coverage level 1 − *α*, the quantile level used during training is

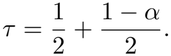

For example, when the target coverage is 0.90, we use *τ* = 0.95.

The pinball loss is defined as

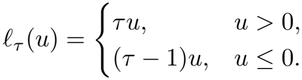

The lower width 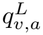 is trained using the signed error *e*_*v,a*_, while the upper width 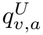 is trained using −*e*_*v,a*_.

The marginal pinball loss is therefore

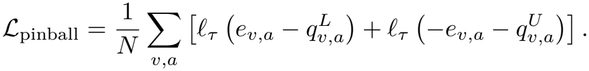

Only using the pinball loss may allow the predicted intervals to become unnecessarily wide. Therefore, we also include a width penalty,

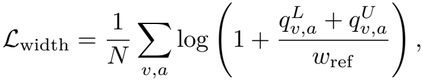

where *w*_ref_ is a reference width.

The final training objective is ℒ= ℒ_pinball_ +*λ*_width_ ℒ_width_. In our implementation, *λ*_width_ = 0.01 and *w*_ref_ = 1.0. The pinball loss helps the model learn the required marginal quantiles, while the width penalty discourages unnecessarily large uncertainty intervals. The gradients from this loss are backpropagated through the three axis-specific heads and the shared GNN backbone. Therefore, the shared representation and the axis-specific uncertainty outputs are learned together.

At this stage, the three axes are still trained marginally, and no joint coverage loss is used.

#### C.4 Conformal-Aware Checkpoint Selection

The training loss is used to optimize the model parameters, but a lower loss does not always mean that the uncertainty intervals have better coverage and smaller volume when we have the final axis width and construct the box to cover the atom. Because of that, we use conformal-aware checkpoint selection to evaluate the model after each training epoch.

At the end of each epoch, the current model is applied to the validation set to obtain the raw uncertainty widths 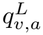 and 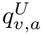. The validation systems are divided into two folds at the system level. For the first fold, the conformal correction is fitted using the second fold and evaluated on the first fold. The procedure is then reversed,

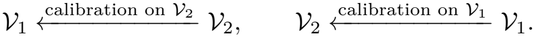

In this way, each validation system is evaluated using a conformal correction fitted without using that system. The calibration and test sets are not used in this checkpoint selection procedure.

For each checkpoint, we evaluate the joint coverage and the size of the three-dimensional uncertainty box. Let

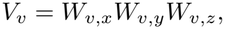

where *W*_*v,a*_ is the calibrated interval width along axis *a*. The efficiency of a checkpoint is measured by

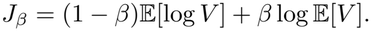

A checkpoint with a small volume can still be unacceptable if its coverage is too low. Therefore, we further adjust the objective using the difference between the target joint coverage and the observed joint coverage,

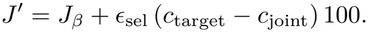

The checkpoint with a smaller value of *J*^*′*^ is preferred, provided that the calibrated intervals remain valid and the joint coverage does not drops lower than the defined safety threshold.

### D Input Features

ConfDock uses 82 input features for each atom. Our predictor uses 63 of these features, with the same subset for all three docking methods. We first remove 17 redundant features that can be reconstructed from other retained columns. We also remove two protein–ligand interface features (columns 74 and 75). Table 7 lists the 19 removed features and their corresponding relationships or removal reasons.

**Table 7:**
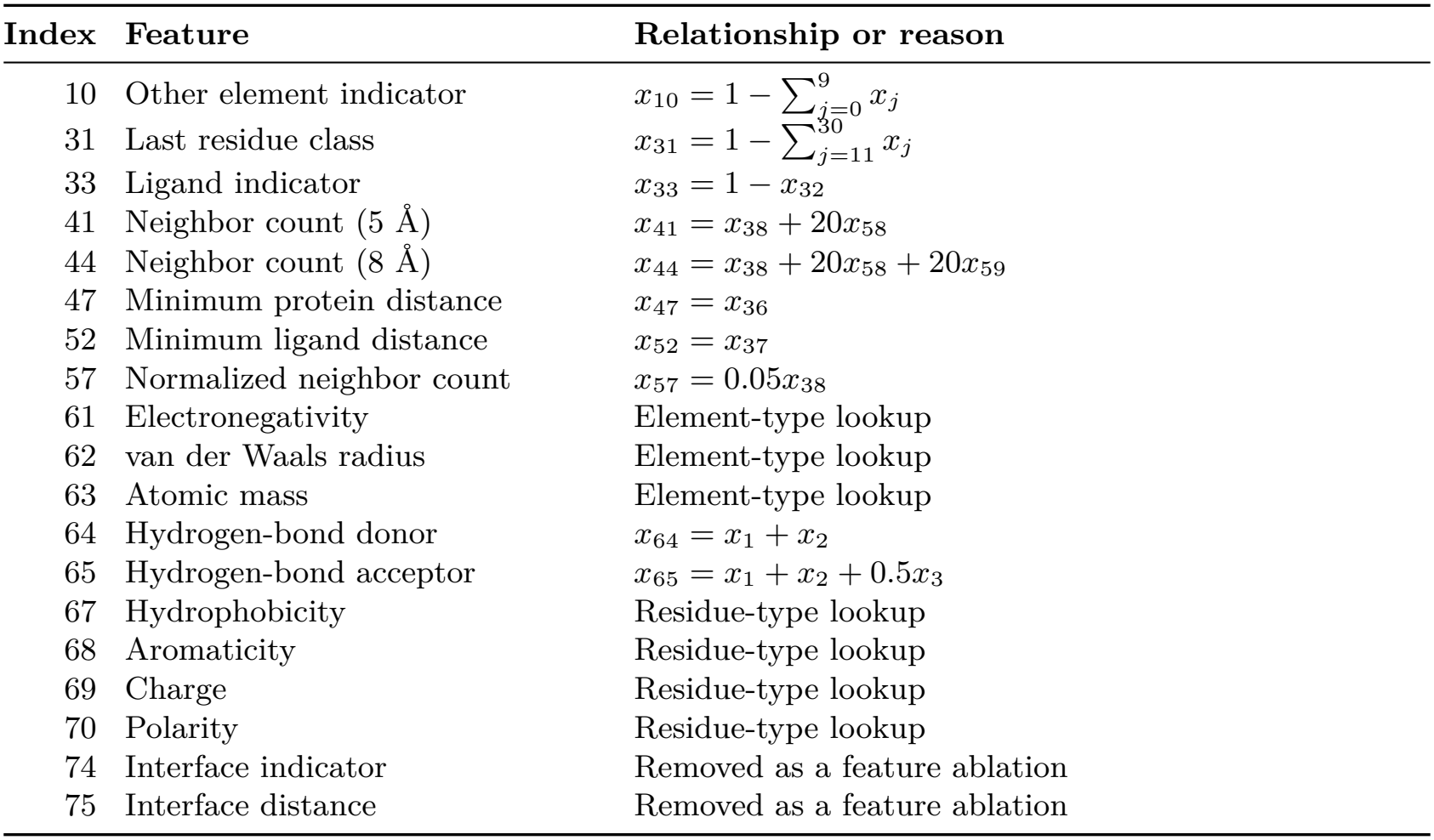
Removed input features from the original 82-dimensional ConfDock feature vector. All indices are zero-based. The first 17 features are redundant, while the last two are removed as a feature ablation.

| **Index** | **Feature** | **Relationship or reason** |
|---|---|---|
| 10 | Other element indicator | $x_{10} = 1 - \sum_{j=0}^{9} x_j$ |
| 31 | Last residue class | $x_{31} = 1 - \sum_{j=11}^{30} x_j$ |
| 33 | Ligand indicator | $x_{33} = 1 - x_{32}$ |
| 41 | Neighbor count (5 Å) | $x_{41} = x_{38} + 20x_{58}$ |
| 44 | Neighbor count (8 Å) | $x_{44} = x_{38} + 20x_{58} + 20x_{59}$ |
| 47 | Minimum protein distance | $x_{47} = x_{36}$ |
| 52 | Minimum ligand distance | $x_{52} = x_{37}$ |
| 57 | Normalized neighbor count | $x_{57} = 0.05x_{38}$ |
| 61 | Electronegativity | Element-type lookup |
| 62 | van der Waals radius | Element-type lookup |
| 63 | Atomic mass | Element-type lookup |
| 64 | Hydrogen-bond donor | $x_{64} = x_1 + x_2$ |
| 65 | Hydrogen-bond acceptor | $x_{65} = x_1 + x_2 + 0.5x_3$ |
| 67 | Hydrophobicity | Residue-type lookup |
| 68 | Aromaticity | Residue-type lookup |
| 69 | Charge | Residue-type lookup |
| 70 | Polarity | Residue-type lookup |
| 74 | Interface indicator | Removed as a feature ablation |
| 75 | Interface distance | Removed as a feature ablation |

The redundancy of the first 17 features was checked using the original atom feature vectors. The direct relationships were verified numerically, while the element- and residue-based lookup features were fully determined by the corresponding one-hot encodings. In particular, columns 41 and 44 contain unnormalised neighbor counts with larger numerical scales, although their information is already represented by the retained columns.

The two interface features are removed separately and are not considered redundant. The resulting 63-dimensional representation retains the other atom-level and local environment features. No new features are added, and the element type used by the atom-type embedding is recovered from the retained element one-hot columns. The feature subset was fixed before the experiments in Section 6 and was not selected using the test systems.

### E Copula Calibration Details

#### E.1 R-vine Copula Calibration

For the R-vine copula, the three-dimensional dependence structure is decomposed into a sequence of pair-copulas. Different pairs can be modeled using different copula families, such as Gaussian, Student-*t*, Clayton, Gumbel, and Frank copulas. This flexibility allows the model to capture nonlinear, asymmetric, and tail dependence among the conformity scores.

Before fitting the copula, the marginal conformity scores are transformed into pseudo-observations on the unit interval:

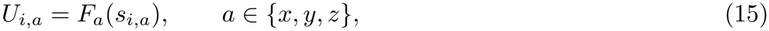

where *F*_*a*_ denotes the empirical marginal CDF of the conformity scores for axis *a* on the calibration set.

Therefore, each calibration sample is represented by the triplet

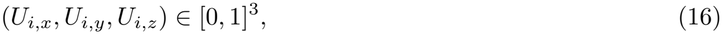

and the R-vine copula is fitted to these transformed conformity scores.

For example, we can choose *X* as the conditioning variable. In this case, a three-dimensional C-vine, which is a special case of an R-vine, can be constructed as follows:

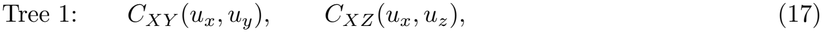

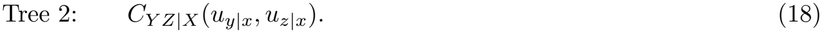

The corresponding joint copula density can then be decomposed as

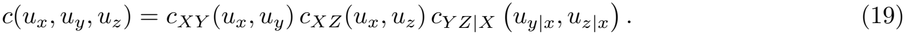

The conditional arguments are obtained through the corresponding *h*-functions:

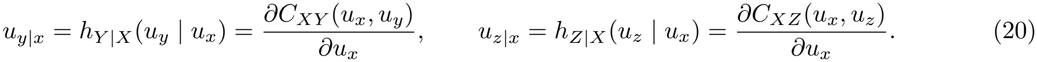

The first tree captures the pairwise dependence between *X* and *Y*, and between *X* and *Z*, while the second tree models the remaining dependence between *Y* and *Z* conditional on *X*.

After fitting the R-vine copula *Ĉ*_vine_, we determine a common copula-level quantile *u*^*\**^ for the target joint coverage 1 − *α* by solving

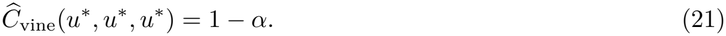

The value *u*^*\**^ is a common probability level in the copula space rather than a calibration threshold in the original conformity-score space. Therefore, it is mapped back to each marginal conformity-score distribution through the corresponding inverse CDF:

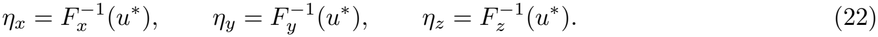

Although the same copula-level quantile *u*^*\**^ is used for all three axes, the resulting calibration thresholds *η*_*x*_, *η*_*y*_, and *η*_*z*_ are generally different because the marginal conformity-score distributions are different.

The calibrated lower and upper uncertainty radii are then defined as

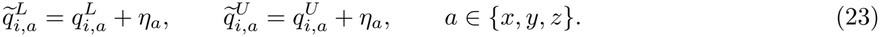

Thus, simultaneous coverage of all three axes is equivalent to

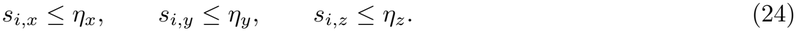

Equivalently, for a signed coordinate error defined as

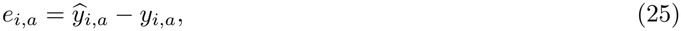

the calibrated coordinate interval along axis *a* is

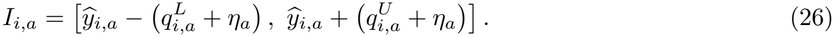

Finally, the three-dimensional uncertainty box is constructed as the Cartesian product of the three calibrated marginal intervals:

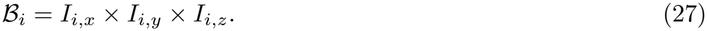

The calibration procedure then targets

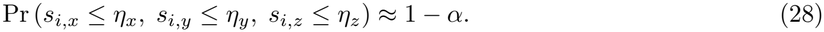

#### E.2 Empirical Copula Calibration

As a nonparametric alternative to the R-vine copula, the empirical copula directly estimates the joint dependence structure from calibration pseudo-observations without assuming a parametric copula family. Given the pseudo-observations *U*_*i*_ = (*U*_*i,x*_, *U*_*i,y*_, *U*_*i,z*_), the empirical copula is defined as

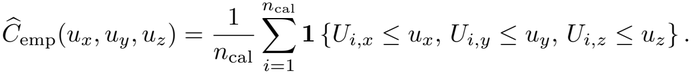

This represents the proportion of calibration samples whose pseudo-observations simultaneously satisfy the three marginal thresholds.

To determine a common threshold for joint coverage, we define

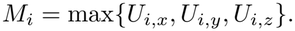

Since the three conditions *U*_*i,x*_ ≤*u, U*_*i,y*_ ≤*u*, and *U*_*i,z*_ ≤*u* are equivalent to *M*_*i*_ ≤*u*, the diagonal empirical copula can be expressed as

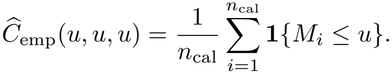

Therefore, the common copula-level quantile for the target joint coverage 1 − *α* is

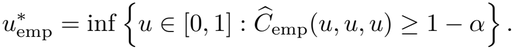

Equivalently, 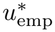 is the empirical (1 − *α*)-quantile of the maximum pseudo-observation scores *M*_*i*_. The corresponding marginal calibration thresholds are obtained using the inverse empirical CDFs:

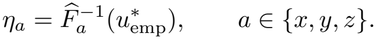

These thresholds are then used to construct the calibrated coordinate intervals and the final three-dimensional prediction box, following the same procedure as the R-vine copula method.

Unlike parametric copula models, the empirical copula requires neither copula-family selection nor correlation-matrix parameterisation. Its dependence structure is determined entirely by the calibration pseudo-observations, making it a useful benchmark for parametric copula calibration. However, empirical estimation may become less stable in high-quantile regions where relatively few calibration samples are available.

#### E.2 Gaussian Copula Calibration

As a parametric approach to dependence-aware calibration, the Gaussian copula models the joint dependence among the three coordinate-wise conformity scores through a correlation matrix. Given the pseudo-observations 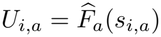, where *a* ∈ {*x, y, z*}, we first apply the inverse standard normal transformation:

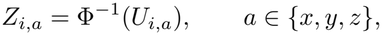

where Φ denotes the standard normal CDF. In practice, rank-based pseudo-observations are used to avoid values of zero or one.

The correlation matrix 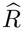 is estimated from the transformed scores (*Z*_*i,x*_, *Z*_*i,y*_, *Z*_*i,z*_). The resulting three-dimensional Gaussian copula is defined as

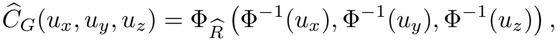

where 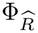 is the trivariate standard normal CDF with correlation matrix 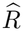.

To determine a common threshold for the target joint coverage 1 − *α*, we solve

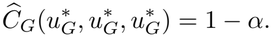

The threshold 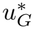 is obtained numerically using the bisection method. The corresponding marginal calibration thresholds are then given by

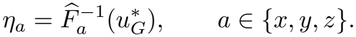

These thresholds are used to construct the calibrated coordinate intervals and the final three-dimensional prediction box. Compared with the R-vine copula, the Gaussian copula is computationally simpler and requires fewer parameters. However, it cannot capture asymptotic tail dependence, which may limit its ability to model simultaneous extreme conformity scores across the three coordinate axes.

### F Density-Ratio Estimation

To evaluate the robustness of our method under distribution shift, we study the difference between the calibration and test distributions. Conformal calibration uses the calibration set to estimate uncertainty for future samples. If the calibration and test distributions are different, the calibration scores may not fully represent the uncertainty of the test samples. We use density ratio estimation (DRE) to adjust the contribution of calibration samples according to their similarity to the test distribution.

For each protein–ligand system *i*, suppose that there are *P*_*i*_ predicted poses. Each pose has a graph-level representation **g**_*i,p*_ ∈ ℝ^256^, obtained from the shared GNN encoder. Since the distribution split is defined at the system level, we aggregate the pose-level representations by taking their mean,

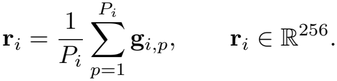

The resulting vector **r**_*i*_ is used as the system-level representation for distribution shift detection.

Directly using all 256 dimensions may lead to unstable domain classification when the number of systems is limited. We first standardize each feature dimension and then apply principal component analysis (PCA) to reduce the representation dimension. Let the reduced representation still be denoted by **r**_*i*_ for simplicity. All preprocessing steps, including standardization and PCA, are fitted only on the training portion of each cross-validation fold and then applied to the corresponding held-out fold.

We define a binary domain label

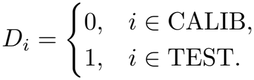

Let *p*_*C*_(**r**) and *p*_*T*_ (**r**) denote the distributions of the system-level representation in the calibration and test sets. The density ratio is defined as

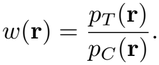

Instead of estimating the two densities separately, we convert the density-ratio estimation problem into a binary classification problem. Let *n*_*C*_ and *n*_*T*_ be the numbers of calibration and test systems. For a system sampled from the combined calibration and test data,

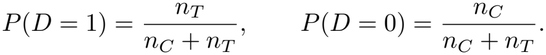

By Bayes’ rule,

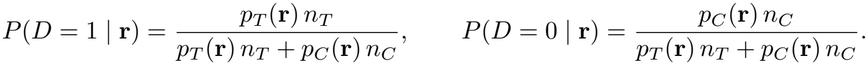

Taking the ratio of these two probabilities gives

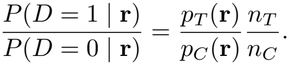

Therefore, the density ratio can be written as

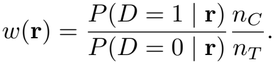

Let *g*(**r**) = *P* (*D* = 1 | **r**). Since *P* (*D* = 0 | **r**) = 1 − *g*(**r**), we obtain

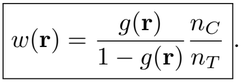

Taking the logarithm,

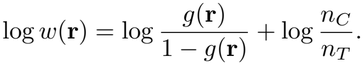

Using the definition of the logit function, logit 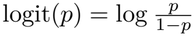, we finally obtain

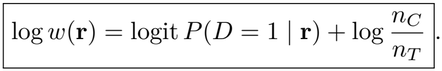

This relation shows that the functional form learned by the domain classifier directly determines the functional form of the estimated log density ratio.

We consider three candidate domain classifiers with different levels of model flexibility. The first model is a linear logistic regression applied to the standardized representation. Its logit has the form

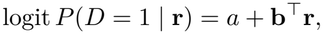

which means that log *w*(**r**) is linear in the representation. This model is simple and less likely to overfit, but it cannot represent nonlinear or interaction-based distribution shifts.

The second model is a second-order polynomial logistic regression. The input representation is expanded to include the original features, squared terms, and pairwise interaction terms,

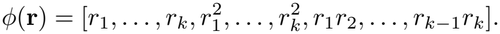

For a *k*-dimensional PCA representation, the expanded feature dimension is

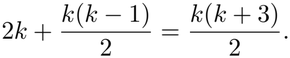

The classifier has the form

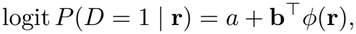

which allows the estimated log *w*(**r**) to contain quadratic and interaction effects. Because polynomial expansion increases the number of parameters, the regularization strength *C* is selected using an inner five-fold cross-validation procedure. Candidate values of *C* are searched over nine logarithmically spaced values from 10^*−*3^ to 10, and the value with the lowest validation log-loss is selected.

The third candidate is a gradient boosting model. We use a histogram-based gradient boosting classifier with tree depth 2, learning rate 0.05, at most 300 boosting iterations, minimum leaf size 30, and *L*_2_ regularization of 1. Early stopping is used during training. A Platt calibration step is applied to improve the calibration of the predicted domain probabilities. Compared with the two logistic models, the boosting model can represent nonlinear and threshold-like changes in the density ratio without requiring a fixed polynomial form.

The three candidate classifiers are selected using stratified five-fold out-of-fold (OOF) log-loss. The combined CALIB and TEST systems are divided into five folds while approximately preserving the proportion of the two domain labels in each fold. For fold *k*, the classifier is trained using the other four folds and predicts the domain probabilities for the held-out fold. This procedure is repeated five times, so every system receives one OOF probability from a model that has not been trained on that system.

For candidate classifier *m*, let *ĝ*_*m*_(**r**_*i*_) denote the OOF estimate of *P* (*D*_*i*_ = 1 | **r**_*i*_). The OOF log-loss is

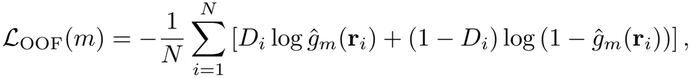

where *N* = *n*_*C*_ + *n*_*T*_ .

The selected classifier is

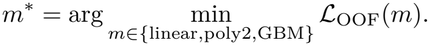

If a classifier predicts 0.5 for every system, its log-loss is − log(0.5) = log 2 ≈ 0.693.

An OOF log-loss below this value indicates that the representation contains information that can distinguish the calibration and test distributions. We use log-loss instead of AUC for classifier selection because the density-ratio weight depends on the numerical value of the predicted probability, not only on its ranking. An overconfident probability close to 0 or 1 can produce an extremely large density ratio even when the classification ranking is reasonable.

After selecting the classifier *m*^*\**^, we directly use its OOF probabilities to compute the density-ratio weights,

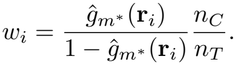

Only the weights corresponding to calibration systems are used in the weighted conformal calibration procedure. We use OOF probabilities instead of probabilities obtained by fitting and evaluating the classifier on the same systems, since in-sample predictions can strongly favor flexible models and generate unstable or overly extreme weights.

The estimated calibration weights are normalized to have mean one,

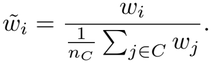

### G Additional Controlled-Shift Experiments

#### G.1 Validation-Based Shift Screening

We first conduct a validation-based screening experiment to identify docking methods for which covariate shifts induce substantial changes in joint coverage. For each docking method, we fix the calibration thresholds and evaluate atom-level coverage on 450 validation systems.

For each system *s*, let *C*_*s*_ denote its empirical atom-level joint coverage and *n*_*s*_ the number of atom-level observations. We simulate distribution shifts by assigning systems to two populations according to

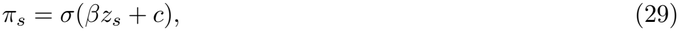

where *π*_*s*_ is the probability of assigning system *s* to the simulated test population, *σ*(*t*) = 1*/*(1 + *e*^*−t*^) is the logistic function, *z*_*s*_ is a rank-normalized system-level covariate, and *β*∈ {−1, −0.5, 0.5, 1} controls the direction and strength of the shift. The intercept *c* is selected so that half of the systems are assigned to the test population in expectation.

The predicted difference in atom-pooled joint coverage between the simulated test and calibration populations is

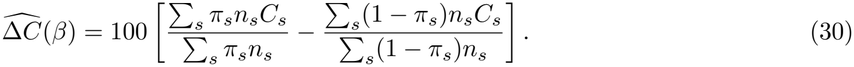

For each docking method, we report the maximum absolute predicted gap across candidate covariates and shift strengths. This screening uses validation outcomes only and does not access held-out test labels.

As shown in Table 8, Protenix exhibits a substantially larger predicted coverage gap (8.83 pp) than AutoDock Vina (1.12 pp) and MedusaGraph (1.31 pp). Only Protenix exceeds the predefined 2-pp screening threshold. We therefore select Protenix for the controlled-shift stress tests.

**Table 8:** Validation-based screening of coverage-relevant distribution shifts. Values indicate the maximum absolute predicted joint-coverage gap in percentage points (pp).

| **Docking Method** | **Max. Gap (pp)** | **Above 2 pp** |
|---|---|---|
| Protenix | **8.83** | Yes |
| AutoDock Vina | 1.12 | No |
| MedusaGraph | 1.31 | No |

#### G.2 Controlled-Shift Construction

We use 899 protein–ligand systems and keep the trained predictor and its predictions fixed. Distribution shifts are introduced by changing the assignment of systems to calibration and test populations, rather than modifying their molecular structures or prediction errors.

For each system *s*, the test-assignment probability is

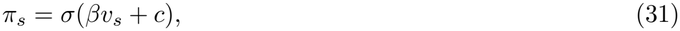

where *v*_*s*_ is a standardized shift-generating covariate, *β* controls shift direction and strength, and *c* balances the expected population sizes. The corresponding oracle density ratio is

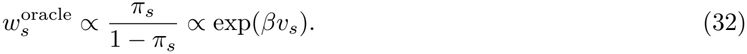

We consider three individual shift directions, PCA01, PCA19, and PCA07, selected through validation-based screening of principal-component scores derived from 256-dimensional system-level GNN representations. Specifically, we evaluate the first 20 principal components on 450 validation systems using the predicted absolute joint-coverage gap under simulated shifts with *β* ∈{−1, +1 }. The components are ranked by their coverage-gap leverage, and the three highest-ranked directions exceeding a 2-percentage-point threshold are selected. This procedure selects PCA01, PCA19, and PCA07, with estimated leverage values of 7.48, 3.84, and 2.49 percentage points, respectively. The selection is based on coverage sensitivity rather than explained variance and does not use held-out test outcomes. This validation-based selection is intended to identify coverage-sensitive directions for controlled stress testing, rather than to characterize the most prevalent distribution shifts in real-world molecular docking. The selection criterion is based on predicted coverage deviations under unweighted calibration, not on the performance of DRE on held-out test systems.

Let *z*_1_, *z*_2_, *z*_3_ denote the normalized scores of these three selected components. We additionally construct three combined directions:

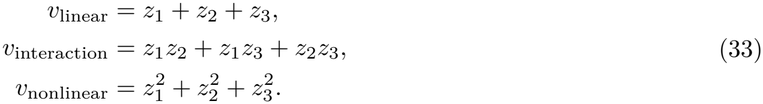

Each combined covariate is standardized before generating the assignments. We evaluate *β* ∈ {−2, −1, −0.5, 0, +0.5, +1, +2 }. The sign of *β* determines the direction of the shift, rather than whether the test distribution is more difficult.

Each nonzero setting is evaluated over 30 repetitions at a target joint coverage of 90%. DRE uses the top 20 principal components of the 256-dimensional graph embeddings. The shift-generating covariates determine the assignment mechanism, while the domain classifier estimates density ratios from the learned representations.

We compare three calibration strategies:

- **Standard:** Unweighted copula-based calibration.
- **Oracle:** Calibration using density ratios derived from the known assignment mechanism.
- **DRE:** Calibration using density ratios estimated by the domain classifier. The signed coverage gap is defined as

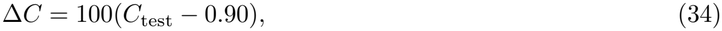

where *C*_test_ is the empirical atom-level joint coverage fraction. Positive values indicate overcoverage, while negative values indicate undercoverage. We report the mean signed gap and mean median prediction-box volume across 30 repetitions for each setting.

#### G.3 Density-Ratio Diagnostics and Null-Shift Behavior

##### Density-Ratio Estimation and Effective Sample Size

As |*β*| increases from 0.5 to 1 and 2, the mean domain-classification AUC rises from approximately 0.57–0.58 to 0.67–0.68 and 0.79, respectively, while the effective sample size (ESS) fraction decreases from 0.82–0.84 to 0.56–0.59 and 0.32. Stronger shifts therefore produce more distinguishable calibration and test distributions but also more variable importance weights. The ESS floor is set to 0.20, and some individual settings approach this limit. Weight stabilization, density-ratio estimation errors, and the use of only 20 of the original 256 embedding dimensions may explain the remaining differences from Oracle calibration.

##### Null-Shift Behavior

At *β* = 0, a separate experiment uses 30 repetitions and 99 permutations per repetition. The permutation-based shift-detection gate activates in 2 of the 30 repetitions (6.7%), close to its nominal 5% significance level. Standard and gated DRE achieve nearly identical test joint coverage (89.84% and 89.83%, respectively) and mean median prediction-box volumes (781.6 and 778.5 Å^3^, respectively). These results suggest that the shift-detection gate generally avoids unnecessary reweighting when no controlled distribution shift is introduced.

### H Joint Calibration Coverage and Validity

Our empirical copula calibration follows the construction of Messoudi et al. (2021). For each calibration atom *i*, let *s*_*i,a*_ denote the nonconformity score along axis *a* ∈ {*x, y, z}* . The scores are transformed using the empirical marginal distributions:

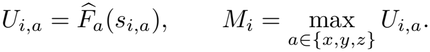

The empirical copula level used in our implementation is defined as

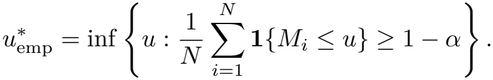

The corresponding marginal thresholds are

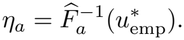

#### Joint calibration coverage

By the definition of *M*_*i*_, the event 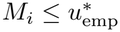 is equivalent to

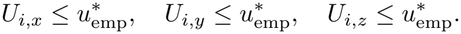

Since the marginal transformations are non-decreasing and the thresholds are obtained from their empirical generalized inverse transformations, these conditions imply

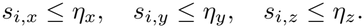

Consequently,

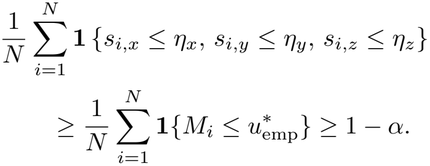

Thus, Empirical Copula achieves at least the target joint coverage on the calibration set. This is an empirical calibration property and does not by itself establish a finite-sample test-coverage guarantee.

#### Finite-sample test coverage

For comparison, consider an idealized split conformal setting in which the marginal score transformations are fixed independently of the calibration data, and the resulting scalar calibration scores *M*_1_, …, *M*_*N*_ and test score *M*_test_ are exchangeable.

Standard split conformal prediction uses the corrected order-statistic index

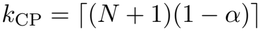

and the corresponding threshold

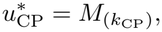

where *M*_(*k*)_ denotes the *k*-th smallest calibration score, with the threshold taken as +∞ when *k*_CP_ *> N* .

Under the stated assumptions, the standard split conformal argument gives

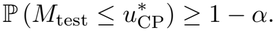

Importantly, this theoretical threshold differs from the empirical threshold 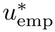 used in our implementation. The latter corresponds to the order-statistic index

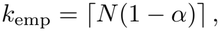

rather than the finite-sample corrected index *k*_CP_.

Therefore, even under the idealized exchangeability assumptions, the empirical threshold does not generally provide the exact nominal finite-sample guarantee.

Furthermore, our implementation estimates the marginal transformations and the joint calibration threshold using the same calibration atoms. Atoms from the same protein–ligand system may also be dependent. Consequently, the assumptions required for the standard split conformal guarantee do not directly hold for the current implementation. We therefore report test joint coverage empirically.

For Gaussian and R-vine copulas, the calibration levels are determined from fitted parametric dependence models. These methods do not directly enforce the empirical joint coverage target and do not provide the same distribution-free finite-sample guarantee. Under distribution shift, DRE further relies on estimated density-ratio weights, whose accuracy also affects the calibration results.

### I Calibration Sensitivity Curve

Figure 4 examines the sensitivity of the calibration methods to different nominal joint coverage levels. Across the three docking methods, empirical copula calibration generally achieves test coverage closer to the nominal level than independence-based calibration, particularly at lower coverage targets.

**Figure 4:**
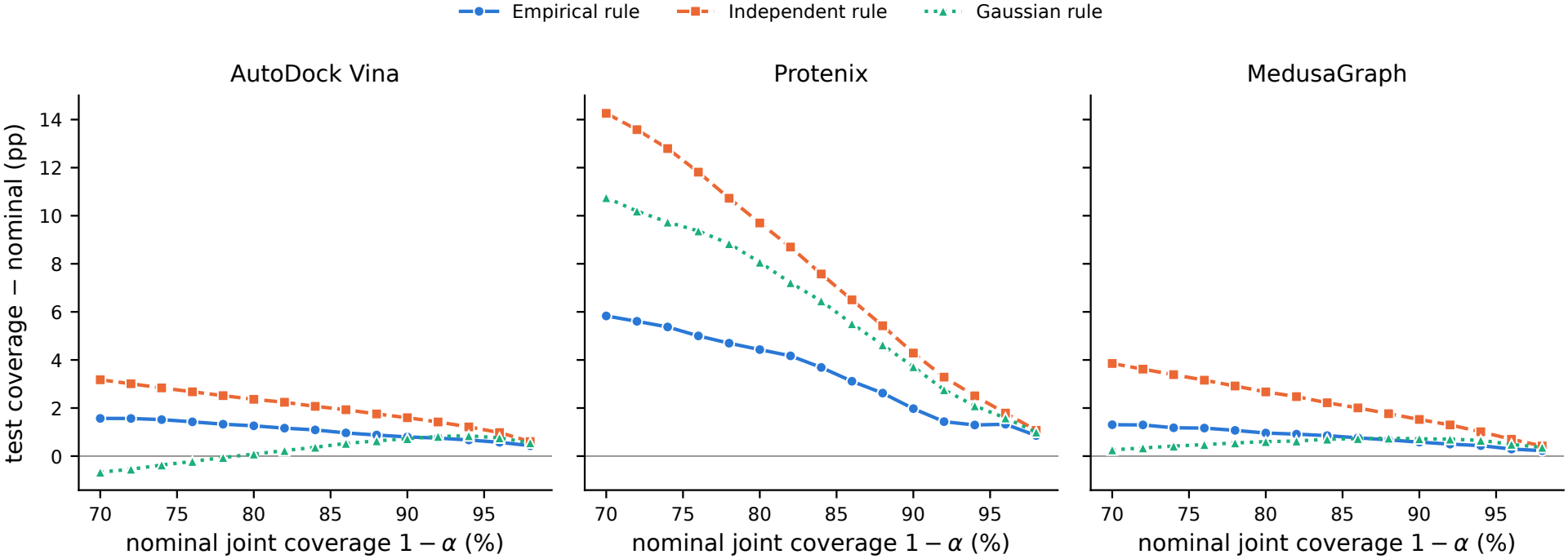
Calibration sensitivity across nominal joint coverage levels for AutoDock Vina, Protenix, and MedusaGraph. The horizontal axis represents nominal joint coverage 1 − *α*, and the vertical axis shows the difference between achieved test joint coverage and nominal coverage, measured in percentage points (pp). Empirical, Independent, and Gaussian calibration rules are compared. The horizontal zero line represents exact agreement with the nominal level. Positive values indicate overcoverage, while negative values indicate undercoverage.

The differences are most pronounced for Protenix, where the Independent and Gaussian rules exhibit substantial overcoverage at lower nominal levels. Empirical copula calibration reduces this conservatism, although some overcoverage remains. For AutoDock Vina and MedusaGraph, the differences between calibration methods are smaller, especially as the nominal coverage approaches 90% and above.

Overall, these results suggest that empirical copula calibration provides a more consistent adjustment for coordinate dependence across a range of nominal coverage levels, rather than only at the 90% target used in the main experiments.

### J Additional Molecular Group Split Analysis

This appendix examines three complementary explanations for the large prediction boxes observed for Protenix under the Molecular Group Split: (i) the distribution of docking errors, (ii) adaptive-width generalization across held-out molecular groups, and (iii) incidental composition differences between the calibration and test subsets. These analyses are diagnostic and do not establish the independent causal contribution of each factor. The Molecular Group Split is constructed using identity-connected groups; calibration and test are assigned symmetrically from the same pool of smaller held-out groups.

#### J.1 Docking Error Distributions and Large-Scale Ligand Misplacement

We first examine differences in docking error distributions between random and Molecular Group Splits to understand why Protenix requires substantially larger prediction boxes under the Molecular Group Split.

##### Changes in the error distribution

Protenix exhibits a pronounced two-mode pattern in its Euclidean ligand-atom displacement distribution under both splits. As summarized in Table 9, the low-error peak changes only slightly, from 0.87 Å under the random split to 1.03 Å under the Molecular Group Split. In contrast, the high-error peak shifts markedly, from 8.71 Å to 16.73 Å. The estimated proportion associated with the high-error mode changes from approximately 32% to 28%. Thus, the most prominent change is in the magnitude of the high-error mode rather than an increase in its estimated mass. For AutoDock Vina and MedusaGraph, the dominant error peaks remain near 5.5 Å under both splits, showing much smaller changes.

**Table 9:** Descriptive peak locations and associated estimated proportions in per-system mean ligand-atom displacement under the random and Molecular Group Splits. For Vina and MedusaGraph, the upper peak contains 95–96% of the estimated mass and represents the distribution’s main body rather than a rare severe-error mode.

| Docking method | Split | Low-error peak | High-error peak |
|---|---|---|---|
| Protenix | Random | 0.87 (68%) | 8.71 (32%) |
| Protenix | Molecular Group | 1.03 (72%) | 16.73 (28%) |
| Vina | Random | 0.70 (5%) | 5.47 (95%) |
| Vina | Molecular Group | 0.71 (4%) | 5.65 (96%) |
| MedusaGraph | Random | 0.75 (5%) | 5.47 (95%) |
| MedusaGraph | Molecular Group | 0.77 (5%) | 5.66 (95%) |

##### Extreme-error prevalence

The shift of the high-error mode is also reflected in the frequency of severe ligand displacements. Using the combined calibration and test populations (899 systems under the random split and 490 under the Molecular Group Split), the proportion of Protenix systems with mean ligand-atom displacement exceeding 10 Å increases from 13.7% to 22.0%. For displacements exceeding 20 Å, the corresponding proportion increases from 4.7% to 10.2%. Despite the slightly smaller estimated mass of the high-error component, this increase is consistent with its shift toward larger displacements. The mode proportions are estimated from the per-system mean ligand-atom displacement of the calibration subset, whereas the exceedance rates above use the combined calibration and test systems.

In contrast, Vina exhibits nearly unchanged proportions of systems exceeding 10 Å (2.6% versus 2.7%), and no systems exceeding 20 Å in either split. The upper tail of the test error distribution also becomes more extreme for Protenix: its 90th percentile increases from 11.8 to 18.8 Å, and its 99th percentile from 24.7 to 29.4 Å. For Vina the corresponding changes are smaller: 7.7 to 8.4 Å (90th percentile) and 10.4 to 11.0 Å (99th percentile).

##### Large-scale ligand displacement

These differences are consistent with the distinct docking procedures used in our experiments. Vina and MedusaGraph operate within predefined binding pockets, constraining the range of possible ligand displacements, whereas Protenix must additionally infer the ligand-binding site. Predictions near the correct binding site may have relatively small coordinate errors, whereas predictions placed in substantially different regions can produce large ligand displacements. Such error regimes offer a plausible explanation for the separated Protenix peaks in Table 9; the peaks do not, by themselves, prove that incorrect pockets were selected.

To characterize severe errors, let **e**_*spv*_ = **ŷ**_*spv*_ − **y**_*spv*_ be the error of ligand atom *v* in pose *p* of system *s*, and define 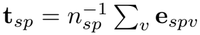, where *n*_*sp*_ is the number of ligand atoms in that pose. The following decomposition holds exactly:

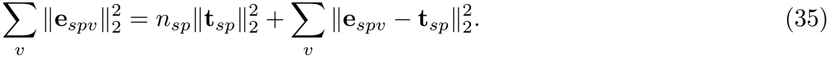

We pool the squared-error contributions across poses within each system and report the fraction attributable to the common-translation term. Among difficult Protenix systems, defined by mean ligand-atom displacement above 10 Å, the median translation shares are 95.7%, 91.9%, 95.5%, and 96.3% on the training, validation, calibration, and test subsets of the Molecular Group Split, respectively. For the remaining systems, the corresponding shares are 17.6%, 22.2%, 21.1%, and 17.7%.

These findings indicate that severe errors contain a dominant coherent displacement component, consistent with substantial ligand misplacement. The decomposition alone does not establish whether an incorrect pocket was selected, nor whether the ligand’s internal geometry was preserved.

##### Implications for calibration

On the original calibration/test partitions, Protenix’s median prediction-box volume under standard empirical-copula calibration increases from 1,168.2 Å^3^ under the random split to 57,644.3 Å^3^ under the Molecular Group Split, an approximately 49-fold increase. We use the original randomsplit partition for this comparison rather than the average over 100 repartitions reported in the random-split main results. The corresponding increases for Vina and MedusaGraph are approximately 1.6-fold.

The larger high-error mode is a plausible source of larger calibration corrections. Its characteristic magnitude nearly doubles under the Molecular Group Split. Correspondingly, the geometric mean of the three axis-specific corrections *η*_*a*_ increases from approximately 4.5 to 17.7 Å, a 3.9-fold increase. The selected empirical-copula level also increases from *u*^*⋆*^ = 0.947 under the random split to 0.957 under the Molecular Group Split. For a fixed nonconformity-score distribution, a higher quantile level cannot decrease the calibration correction; however, the separate contributions of the score distribution and the selected copula level cannot be determined from this comparison.

The raw Euclidean-error distribution does not determine *η*_*a*_ by itself: *η*_*a*_ is a quantile of the coordinate-wise nonconformity scores after accounting for the predicted adaptive widths. Because *η*_*a*_ is applied globally to all atoms, large corrections also enlarge the prediction boxes of systems with relatively small docking errors. Thus, the more extreme docking errors help explain the large Protenix boxes, while the effectiveness of width allocation on systems from held-out molecular groups remains a separate question, examined next.

#### J.2 Training–Held-out Shift and Adaptive-Width Generalization

We next investigate the generalization of adaptive uncertainty widths from training systems to Protenix systems from held-out molecular groups. We examine differences in prediction difficulty, controlled tail oversampling, and representation-space novelty. These diagnostics characterize the observed generalization gap but do not establish its unique causal explanation.

##### Training–held-out distributional differences

The Molecular Group Split separates training, validation, calibration, and test systems by molecular groups. The seven largest groups are assigned to training, the next ten to validation, and the remaining smaller groups to calibration and test. The largest training group alone contains 1,674 of the 1,998 training systems. Consequently, the training and held-out subsets differ not only in group identity but also potentially in their molecular compositions and difficulty distributions.

For Protenix, systems with mean ligand-atom displacement exceeding 10 Å account for 10.4% of the training set, compared with 11.7%, 21.9%, and 22.2% of the validation, calibration, and test sets, respectively. The calibration set therefore contains approximately twice the proportion of difficult systems observed during training.

To examine the effectiveness of adaptive uncertainty prediction, we define the tail-width ratio as the mean predicted half-width among the atoms with the largest 5% of absolute coordinate errors, divided by the overall mean predicted half-width, averaged across the three coordinate axes. Larger values indicate that relatively more uncertainty is assigned to high-error atoms.

As shown in Table 10, the tail-width ratio decreases from 5.16 on training systems to 2.18 on validation systems, and further to 1.84 and 1.80 on calibration and test systems. The system-level Spearman correlation between mean predicted width and mean absolute error also decreases substantially on the held-out subsets.

**Table 10:** Adaptive-width diagnostics for Protenix under the Molecular Group Split. Hard systems have mean ligand-atom displacement exceeding 10 Å. The tail-width ratio measures the relative predicted half-width assigned to the largest 5% of coordinate errors. The correlation is computed between system-level mean predicted half-width and mean absolute coordinate error. Training values are in-sample for the width predictor.

| Subset | Systems | Hard (%) | Tail-width ratio | Spearman ρ |
|---|---|---|---|---|
| Train | 1998 | 10.4 | 5.164 | 0.854 |
| Validation | 206 | 11.7 | 2.182 | 0.470 |
| Calibration | 247 | 21.9 | 1.836 | 0.525 |
| Test | 243 | 22.2 | 1.801 | 0.486 |

Importantly, validation systems have a difficulty prevalence close to that of training systems, yet their tail-width ratio is already substantially lower. Approximately 90% of the observed train-to-calibration decline in this ratio occurs by the validation subset, before the increase in difficult-system prevalence seen in calibration.

Thus, the deterioration cannot be explained solely by the larger proportion of difficult systems in calibration. It is already apparent on systems belonging to molecular groups excluded from training. Nevertheless, training and validation also differ in other characteristics, including group identity and group size. These comparisons therefore cannot isolate the effect of group novelty from other distributional differences.

Some decline from training to held-out systems is expected for any predictor, because the training values are in-sample. The random split provides a reference without group exclusion: there, Protenix’s calibration tail-width ratio retains 62% of its training value, compared with 36% under the Molecular Group Split, whereas Vina and MedusaGraph retain 88–92% under both splits. The additional loss is therefore concentrated in Protenix under the Molecular Group Split.

##### Tail-oversampling intervention

To examine whether the underrepresentation of difficult systems during training explains the width deterioration, we conduct a controlled tail-oversampling experiment. Difficult training systems are identified by their mean ligand-atom displacement, and their sampling weight is doubled, which raises their share of sampled training graphs from 9.7% to 17.7%, approaching the prevalence of difficult systems in calibration. Both arms are retrained from scratch with a shortened budget of at most 40 epochs (minimum 10, patience 10) and differ only in the sampler; the architecture, split, seed, and all other settings are shared. The control therefore differs slightly from the original long-budget model used for the diagnostics in Table 10; the two runs are not directly comparable in their absolute metrics.

On the calibration subset, the tail-width ratio changes from 1.784 in the control to 1.820 after oversampling, an increase of approximately 2.0%. We also report the geometric mean over the three axes of 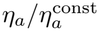, where 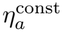 is the calibration correction obtained when every atom receives the mean predicted widths of its axis. A value of 1 means the global corrections coincide for these two width assignments; it does not by itself establish that adaptive widths provide no benefit. Smaller values indicate that the adaptive widths reduce the global correction more. This ratio increases from 0.993 to 1.023, the opposite of the improvement in the tail-width ratio.

The two primary diagnostics therefore show small changes in inconsistent directions. Since this experiment uses a single seed and the selected checkpoints occur at different epochs (6 and 4), these changes cannot be distinguished reliably from training variability. We find no detectable recovery of adaptive tail-width allocation under this intervention.

This result does not exclude broader training–held-out distribution shift as a contributing factor. Doubling the frequency of existing difficult training systems does not introduce new molecular groups or ensure generalization to unfamiliar molecular environments. It therefore tests a specific training-frequency hypothesis rather than the full range of possible distributional differences.

##### Representation-space novelty

We further examine whether adaptive-width under-allocation increases continuously with distance from the training distribution. Using system-level embeddings from the trained GNN, we measure each held-out system’s mean distance to its ten nearest training systems.

For each training system, we compute the same quantity using its ten nearest other training systems, excluding the system itself; the median is 11.67. For validation and calibration systems, the median distances to their ten nearest training systems are 15.08 and 15.73, respectively. Thus, validation and calibration have similar distances to the training sample under this representation. Within calibration, however, the Spearman correlation between embedding distance and the width residual, defined as the residual of a regression of log mean predicted width on log mean error across systems, is positive (*ρ* = 0.402, 95% system-bootstrap interval [0.277, 0.518]), rather than the negative association expected if greater embedding distance systematically produced greater width under-allocation. The interval resamples systems and may be too narrow when systems from the same molecular group are correlated.

These results do not support a simple monotonic distance-based explanation of the generalization gap. However, embedding distance is only a proxy for biological novelty, and the system-level diagnostic may not capture failures concentrated in individual high-error atoms. Broader training–held-out distributional differences therefore remain a possible explanation.

##### Interpretation and limitations

Together, these diagnostics indicate reduced adaptive-width effectiveness on Protenix systems from held-out molecular groups. The decline is already evident in validation, where the prevalence of difficult systems remains close to that of training, and cannot be attributed solely to the greater frequency of difficult systems in calibration.

The tail-oversampling experiment does not reveal a clear recovery of width adaptivity, while the representation-space analysis does not support a simple monotonic relation between embedding distance and width under-allocation. These results do not rule out broader differences in molecular composition, group identity, or prediction difficulty between training and held-out subsets. Moreover, the training diagnostics are in-sample, and the oversampling comparison is limited to a single seed and shortened training budget. The relative contributions of these factors therefore remain unresolved.

Improving adaptive-width generalization across unseen molecular groups is a promising direction for future work. The dependence-aware copula calibration in our framework addresses coordinate-wise score dependence, which is distinct from the generalization of the underlying width predictor.

#### J.3 Calibration–Test Composition Imbalance and DRE Sensitivity

The calibration and test subsets are symmetric group-level partitions of the same held-out pool, but their realized composition need not match. For Protenix, the fraction of systems with mean Euclidean displacement above 20 Å is 12.6% in calibration versus 7.8% in test, even though the broader 10 Å difficulty shares are similar (21.9% and 22.2%). This difference in the extreme-error tail is consistent with the observed overcoverage under standard calibration; it does not, on its own, prove that the sample imbalance caused the coverage deviation. The corresponding DRE analysis adjusts the empirical-copula joint calibration for differences in the observed system-level representation distribution, while leaving marginal score transformations unweighted.

As shown in Table 11, DRE produces test joint coverage between 89.91% and 90.04% for *k* ∈ {40, 50, 60, 70}, compared with 92.31% under standard calibration. At *k* = 50, median volume decreases from 57644.30 to 29271.04 Å^3^ and mean volume from 75977.14 to 42581.63 Å^3^. The OOF domain-classification AUC rises from 0.686 to 0.717 across the PCA settings, whereas the ESS fraction remains low (0.274–0.292), indicating potentially variable importance weights.

**Table 11:** PCA sensitivity of DRE-weighted empirical-copula calibration for Protenix under the Molecular Group Split. ESS is the effective sample size fraction; prediction-box volumes are in Å^3^.

| *k* | OOF AUC | ESS | Test joint (%) | Median vol. | Mean vol. |
|---|---|---|---|---|---|
| 40 | 0.686 | 0.292 | 89.91 | 27442.78 | 40374.82 |
| 50 | 0.698 | 0.280 | 90.04 | 29271.04 | 42581.63 |
| 60 | 0.714 | 0.274 | 89.95 | 27903.04 | 40931.50 |
| 70 | 0.717 | 0.274 | 90.02 | 28861.41 | 42087.76 |

These results suggest that DRE can improve empirical calibration efficiency by adapting the joint calibration threshold to differences in the realized test composition. However, DRE does not directly address the adaptive-width generalization gap identified in Appendix J.2, and it does not provide an exact finite-sample joint-coverage guarantee under the group-dependent Molecular Group Split.

### K Additional Analysis of Weight Placement

#### Calibration Strategies

Let *s*_*i,a*_ denote the nonconformity score of calibration observation *i* along coordinate axis *a* ∈ {*x, y, z }*, and let *ω*_*i*_ denote its density-ratio weight inherited from the corresponding protein–ligand system. For *N* calibration observations, define the unweighted and weighted marginal score transformations as

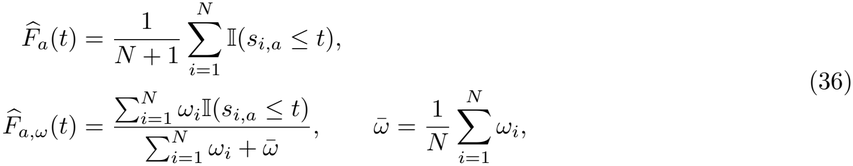

where I(·) is the indicator function.

For a target joint coverage level 1 − *α*, the two strategies are defined as

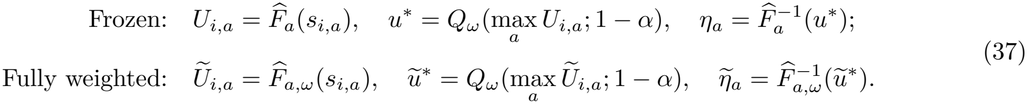

Here *Q*_*ω*_( ; 1 − *α*) denotes the weighted (1 − *α*)-quantile of the transformed joint scores. Both strategies use the same importance weights and empirical copula. The distinction is whether the marginal score transformations are also weighted.

#### Experimental Details

We evaluate six controlled shift directions under *β* ∈ {±0.5, ±1, ±2} using Protenix. Each experiment is repeated 30 times at 90% target joint coverage. We compare exact oracle weights exp(*βv*_*s*_) without stabilization and stabilized DRE weights estimated using PCA-20 graph embeddings or three selected system-level features.

#### Additional Results

Under moderate shifts (|*β*| ≤1), frozen and fully weighted marginals exhibit similar coverage, with differences within 0.03 pp. For positive moderate shifts, fully weighted calibration produces larger prediction boxes in all 12 oracle settings and 23 of 24 DRE settings, with average volume increases of approximately 1–4%.

Under strong shifts (| *β*| = 2), the differences become more variable, especially with unstabilized oracle weights, whose effective sample size fraction is approximately 0.14–0.16. At *β* = −2, the volume ratio ranges from 0.58 to 1.66 across scenarios. At *β* = +2, fully weighted calibration reduces the mean volume ratio to 0.816 but increases absolute coverage deviation from 0.90 to 1.21 pp. Stabilized DRE weights produce smaller differences.

#### Interpretation

When the weighted and unweighted marginal transformations differ across coordinates, fully weighted calibration changes the relative ordering of the transformed scores and therefore the maximum score used for joint calibration. These changes can alter the axis-specific thresholds, potentially affecting both joint coverage and prediction-box volume. The empirical results do not establish a universal advantage for either weighting strategy, but support frozen marginals as a simpler and more stable choice in the evaluated settings.

### L Analysis of Prediction Error Distributions

We compare the marginal and system-difficulty-stratified distributions of ligand-atom prediction errors for three docking methods (AutoDock Vina, Protenix, and MedusaGraph) on the HiQBind systems.

Figure 6 shows the signed coordinate errors along the *x, y*, and *z* axes, while Figure 5 shows the Euclidean error magnitudes. In each panel, the upper plot shows the kernel density estimate across all ligand atoms, while the lower plot divides the atoms into five groups based on a leave-one-pose-out system-level difficulty score (bin 0 easiest, bin 4 hardest). This grouping is used only for retrospective analysis and is not included as an input during testing.

**Figure 5:**
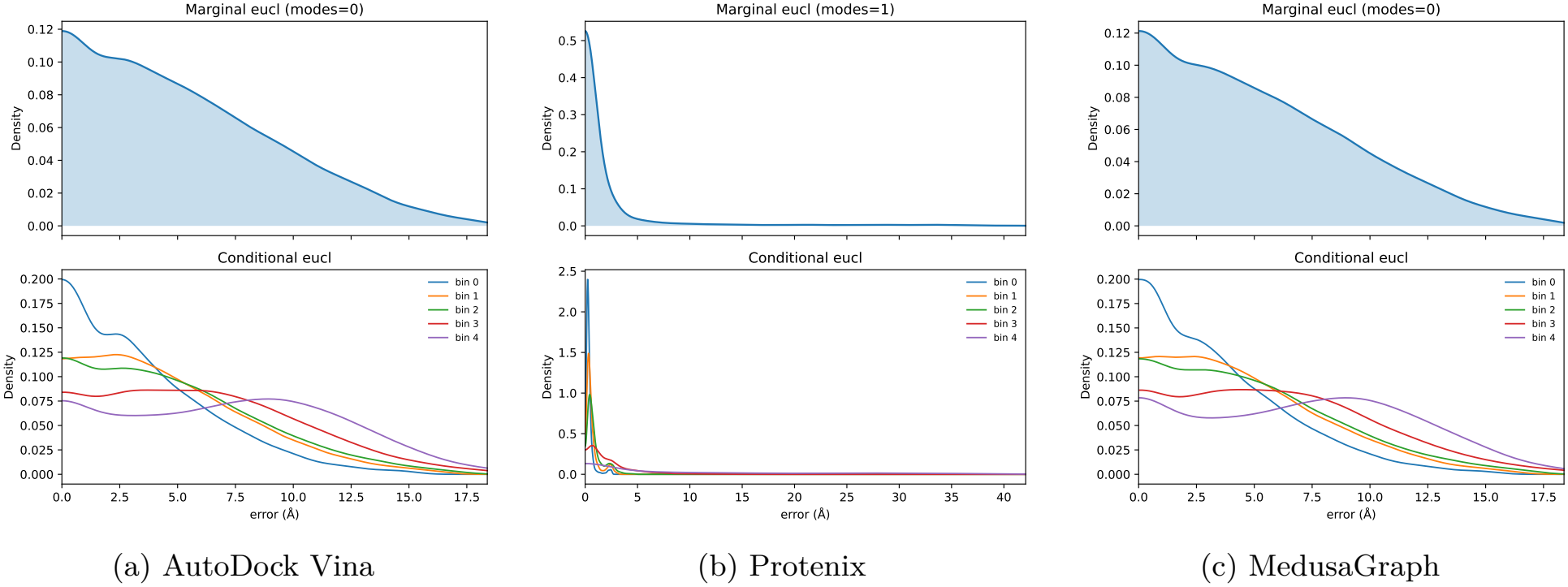
Pooled (upper) and system-difficulty-stratified (lower) kernel density estimates of Euclidean error magnitudes for AutoDock Vina, Protenix, and MedusaGraph. Difficulty bins range from 0 (easiest) to 4 (hardest). Each density is normalized separately; horizontal scales differ across methods.

**Figure 6:**
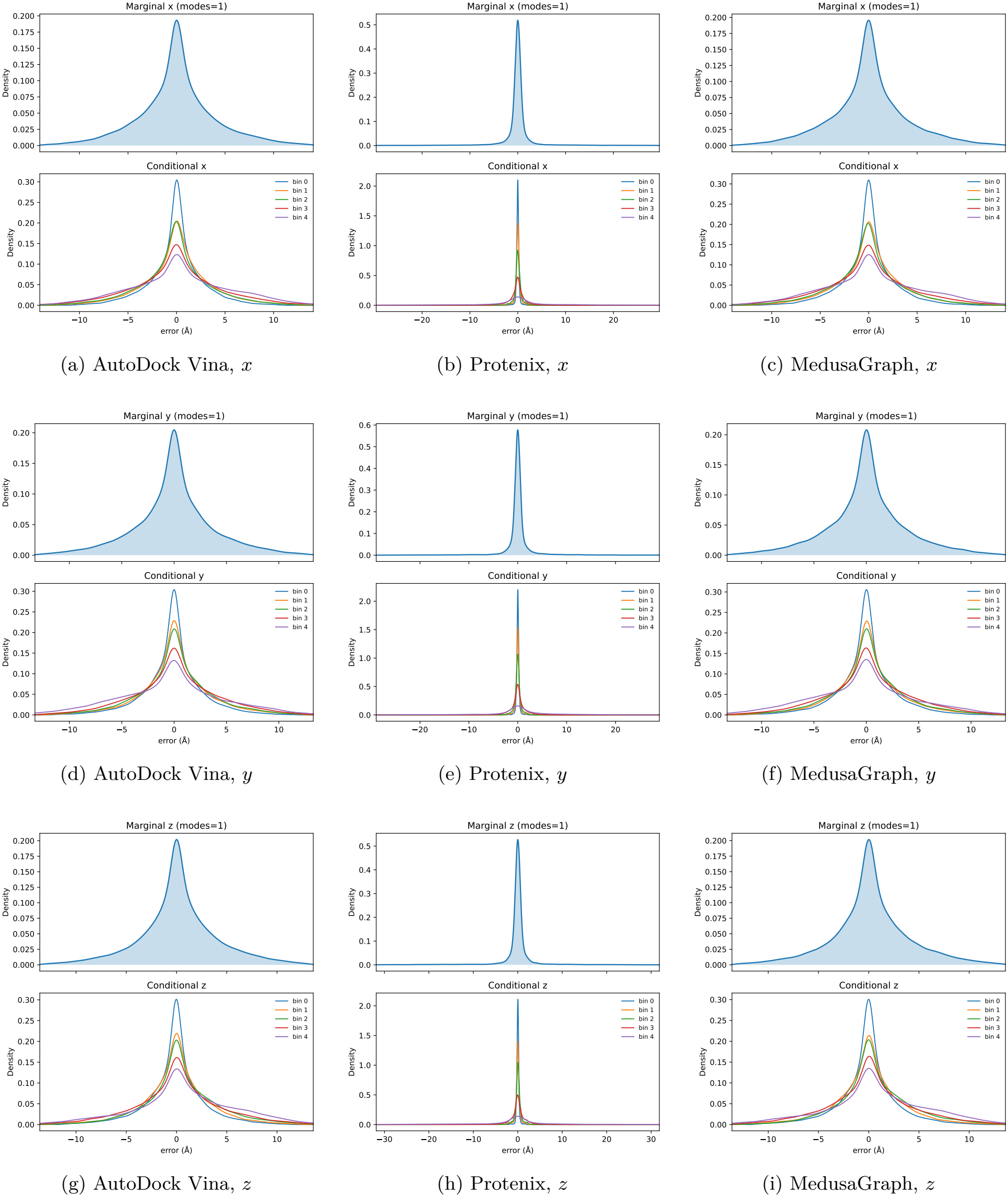
Pooled (upper) and system-difficulty-stratified (lower) kernel density estimates of signed coordinate errors. Rows correspond to the *x, y*, and *z* axes, and columns correspond to AutoDock Vina, Protenix, and MedusaGraph. Difficulty bins range from 0 (easiest) to 4 (hardest). Each density is normalized separately; horizontal scales differ across methods.

Two main observations can be made. First, the error distributions vary across system-difficulty bins. In particular, the Euclidean error distributions for AutoDock Vina and MedusaGraph generally shift toward larger error magnitudes as system difficulty increases, indicating substantial system-level heterogeneity in prediction errors. This motivates the use of a graph-level embedding **g** in each atom representation to capture structural context associated with prediction uncertainty.

Second, the signed error distributions along the *x, y*, and *z* axes exhibit broadly similar marginal patterns within each docking method. However, these marginal distributions alone do not establish statistical dependence between coordinate errors. Since the three coordinate errors jointly determine the spatial prediction error, their dependence structure is relevant when constructing joint three-dimensional prediction regions. This motivates the use of copula-based calibration to account for cross-coordinate dependence.

These density plots provide a descriptive analysis of prediction error distributions rather than a direct evaluation of calibration performance. Each curve is normalized separately, and the horizontal scales differ across methods and error types.

